# Striatonigrostriatal Circuit Architecture for Disinhibition of Dopamine Signaling

**DOI:** 10.1101/2021.06.22.449416

**Authors:** Priscilla Ambrosi, Talia N. Lerner

## Abstract

The basal ganglia operate largely in closed parallel loops, including an associative circuit for goal-directed behavior originating from the dorsomedial striatum (DMS) and a somatosensory circuit important for habit formation originating from the dorsolateral striatum (DLS). An exception to this parallel circuit organization has been proposed to explain how information might be transferred between striatal subregions, for example from DMS to DLS during habit formation. The “ascending spiral hypothesis” proposes that DMS disinhibits dopamine signaling in DLS through a tri-synaptic, open-loop striato-nigro-striatal circuit. Here, we used transsynaptic and intersectional genetic tools to investigate both closed- and open-loop striato-nigro-striatal circuits. We found strong evidence for closed loops, which would allow striatal subregions to regulate their own dopamine release. We also found evidence for functional synapses in open loops. However, these synapses were unable to modulate tonic dopamine neuron firing, questioning the prominence of their role in mediating crosstalk between striatal subregions.

## INTRODUCTION

The striatum is well-known for its roles in motor control and reinforcement learning. The dorsomedial striatum (DMS) is thought to be involved in goal-directed learning, while the dorsolateral striatum (DLS) is thought to be involved in motor skill acquisition and habit formation (Lipton et al., 2019; Yin and Knowlton, 2006). As animals are overtrained in a motor skill task (e.g., accelerating rotarod) or in an instrumental task designed to elicit habit (e.g., random interval training), their behavior becomes more stereotyped and less flexible, and dependence of the behavior shifts from DMS to DLS (Corbit et al., 2012; Derusso et al., 2010; Gremel and Costa, 2013; Sommer et al., 2014; Thorn et al., 2010; Yin et al., 2004, 2005a, 2005b, 2006, 2009).

Both DMS and DLS are richly innervated by dopamine (DA) neurons from the substantia nigra pars compacta (SNc). Although DA axonal fields in striatum are broad (Matsuda et al., 2009), there is topography within the nigro-striatal system that can allow for separate control of DA release in DMS and DLS (Farassat et al., 2019; Ikemoto, 2007; Joel and Weiner, 2000; Lerner et al., 2015). Indeed, DA neurons projecting to DMS and those projecting to DLS display distinct *in vivo* activity patterns (Brown et al., 2011; Hamid et al., 2021; Lerner et al., 2015; Seiler et al., 2020; Tsutsui-Kimura et al., 2020).

How distinct activity in DMS-projecting and DLS-projecting DA neurons arises is a key question. One possibility is that these cells receive distinct inputs (Lerner et al., 2015). In particular, it has been widely hypothesized that DMS-DLS transitions observed during habit formation are regulated by an input circuit to DLS-projecting DA neurons termed the “ascending spiral” (Haber et al., 2000; Lerner, 2020; Lüscher et al., 2020; Yin and Knowlton, 2006). The premise of the ascending spiral hypothesis is that DMS and DLS are connected by a tri-synaptic circuit involving GABAergic neurons in substantia nigra pars reticulata (SNr) and DA neurons in SNc. More specifically, DA neurons are thought to be under tonic inhibition from GABAergic neurons in SNr; spiny projection neurons (SPNs) from DMS can inhibit these SNr GABA cells, and thus disinhibit DLS-projecting DA neurons, allowing for DA release in DLS. The individual steps in this polysynaptic circuit (DMS→SNr, SNr→SNc, and SNc→DLS) are well-established (Chevalier et al., 1985; Freeze et al., 2013; Tepper and Lee, 2007; Tepper et al., 1995). However, it is not necessarily the case that the individual connections link into a continuous polysynaptic circuit (DMS→SNr→SNc→DLS). Indeed, anatomical and electrophysiological work in other basal ganglia circuits supports a largely parallel organization of DMS and DLS subcircuits (Alexander et al., 1986; Lee et al., 2020; Mandelbaum et al., 2019). The idea that an ascending spiral through the midbrain DA system could be a major route of crosstalk between otherwise parallel circuits has been appealing to behavioral neuroscientists, but evidence of a functional circuit at the synaptic level is lacking.

Evidence for the ascending spiral hypothesis stems primarily from anatomical work done in non-human primates (Haber et al., 2000). Following the injection of retrograde and anterograde tracers in striatum, Haber and colleagues uncovered a medio-lateral organization of striato-nigro-striatal circuits. Namely, axon terminals from medial striatum are medially located in SN and overlap with the cell bodies of neurons that project to medial and lateral striatum. Axons from lateral striatum, on the other hand, are laterally located in SN and overlap with cells that project to lateral, but not medial striatum. Thus, there is a proposed asymmetry in which medial striatum could influence DA release in lateral striatum, but lateral striatum would not influence DA release in medial striatum. Critically, however, the overlap of axon terminals and cell bodies is neither necessary nor sufficient for the existence of a functional circuit, especially a polysynaptic circuit requiring an intermediary connection between the axon terminals and cell bodies in question. Therefore, despite its appeal, the ascending spiral hypothesis rests on weak evidence.

A direct test of the tri-synaptic circuit proposed by the ascending spiral hypothesis has been lacking in part because of technological limitations that prevented selective targeting of projection-specific circuit components. We took advantage of recent developments in transsynaptic tracing (Zingg et al., 2017, 2020) and intersectional genetics (Fenno et al., 2014; Poulin et al., 2018) to solve this problem. Then, using slice electrophysiology and optogenetics, we found the first functional evidence supporting the existence of an ascending spiral circuit. Unexpectedly, we also found similar evidence for a descending spiral circuit rather than the asymmetry predicted from anatomy alone. Finally, although we found evidence for synaptic connectivity in spiral circuits, these circuits were not able to modulate the tonic firing of DA neurons. In contrast, closed loop circuits (DMS→SNr→SNc→DMS and DLS→SNr→SNc→DLS) were. These data provide the first thorough electrophysiological description of functional striato-nigro-striatal circuits in naïve mice. While they support the possibility of an ascending spiral circuit, they overall question the prominence of its role in modulating DA activity patterns. How these striato-nigro-striatal circuits operate *in vivo*, especially during habit formation, and why there is a dissociation between synaptic connectivity and the control of DA cell firing are important questions for future investigation. Gaining a better understanding of striato-nigro-striatal circuit organization and function will inform thinking about the mechanisms underlying habit formation and other dopamine- and striatal-dependent behaviors.

## RESULTS

### DLS- and DMS-Projecting Dopamine Neurons Are Robustly Inhibited by SNr

Motivated by the desire to test whether there is a synaptic basis for the ascending spiral hypothesis, and to understand the organization of disinhibitory striato-nigro-striatal circuits more generally, we designed a series of experiments using synaptic physiology in combination with carefully targeted optogenetic stimulation. We began our study by assessing the connectivity of GABAergic SNr cells to DLS- and DMS-projecting DA neurons. Although SNr is a well-known source of inhibitory input onto SNc DA neurons in general (Tepper and Lee, 2007; Tepper et al., 1995), it was unclear whether the likelihood of receiving GABAergic inputs varied depending on the downstream projection target of the DA neuron. Using acute slice electrophysiology, we recorded the responses of projection-defined DA neurons while optogenetically stimulating GABAergic cells in SNr.

We labeled projection-defined DA neurons by injecting red retrobeads into DLS (Figure 1) or DMS (Figure 2). These fluorescently labeled latex beads travel retrogradely from axon terminals to cell bodies and allow for targeted patching of DLS- or DMS-projecting DA neurons in midbrain slices. We are confident that bead-labeled cells are dopaminergic given that (1) bead-labeled cells in SNc were previously shown to be TH+ (Lerner et al., 2015) and (2) all bead-labeled cells recorded in a loose seal configuration in this study (143/143 cells from 24 mice) had wide action potential waveforms (total duration > 2 ms) characteristic of DA neurons (Grace and Bunney, 1983).

**Figure 1.**
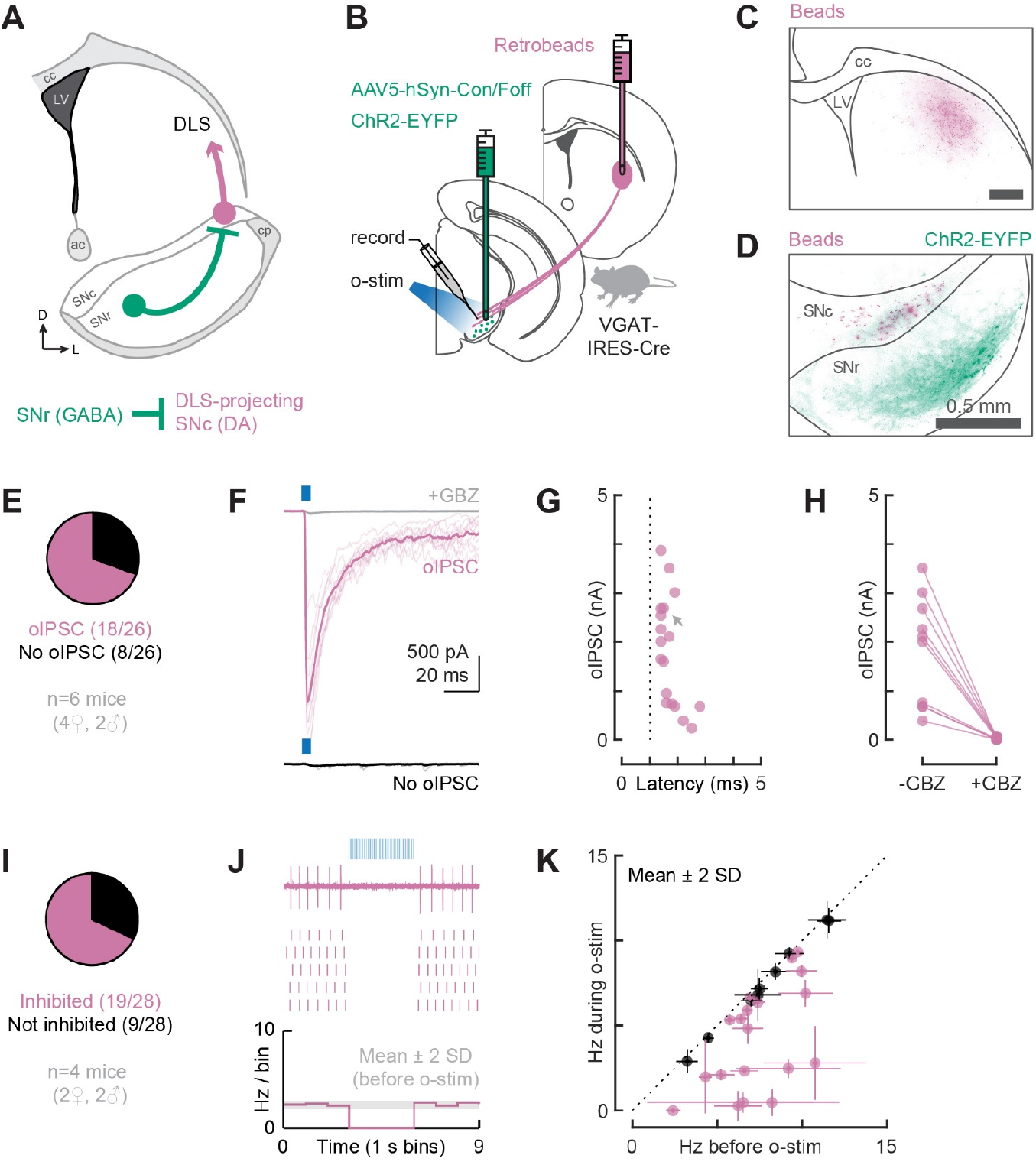
VGAT+ cells in SNr monosynaptically inhibit DLS-projecting DA neurons in SNc and suppress their tonic firing. **(A)** Schematic of the tested circuit. Anatomical landmarks: corpus callosum (cc), lateral ventricle (LV), anterior commissure (ac), cerebral peduncle (cp). **(B)** Experimental design for probing the connection between VGAT+ cells in SNr and DLS-projecting DA neurons in SNc. All injections were done in adult VGAT-IRES-Cre mice. AAV5-hSyn-Con/Foff-ChR2-EYFP was injected into SNr to deliver the excitatory opsin ChR2 to VGAT+ cells. Foff is irrelevant in this setup. Retrobeads were injected into DLS to label the soma of DLS-projecting neurons for targeted patching in midbrain slices. Optogenetic stimulation (o-stim) was delivered via the objective (460 nm, ~20 mW/mm^2^). **(C)** Distribution of retrobeads (magenta) in a representative striatum slice. Scale bar: 0.5mm. **(D)** Distribution of bead-labeled somas (magenta) and ChR2-EYFP-labeled neuropil (green) in a representative midbrain slice. SNc was outlined based on TH immunolabeling. **(E)** Proportion of DLS-projecting neurons recorded in whole-cell mode that did (magenta) or did not (black) respond to the o-stim (5ms pulse) with an optogenetically-evoked inhibitory postsynaptic current (oIPSC). **(F)** Example recording from a cell that did (magenta) or did not (black) show an oIPSC. The oIPSC was absent after gabazine (GBZ) perfusion (gray). Thin lines: individual sweeps. Thick lines: average. **(G)** oIPSC amplitude and onset latency for all responding cells (dotted line = 1ms). A gray arrow indicates the oIPSC shown in F. **(H)** oIPSC amplitude before and after GBZ perfusion for all tested cells. **(I)** Proportion of DLS-projecting neurons recorded in loose seal mode that did (magenta) or did not (black) have their tonic firing suppressed by the o-stim (5ms pulses at 20Hz for 3s). **(J)** Example loose seal recording from a cell that was inhibited by the o-stim. Top: raw data from a single sweep. Middle: raster plot showing action potentials from 5 sweeps. Bottom: histogram of the average firing rate across all sweeps. The gray shaded area indicates mean±2SD for the baseline firing rate. Note that the firing rate drops below mean-2SD during the o-stim. **(K)** Average firing rate before and during the o-stim for all cells recorded in loose seal mode (inhibited cells: magenta; not inhibited: black). Error bars represent ±2SD. Dotted line: unity.

**Figure 2.**
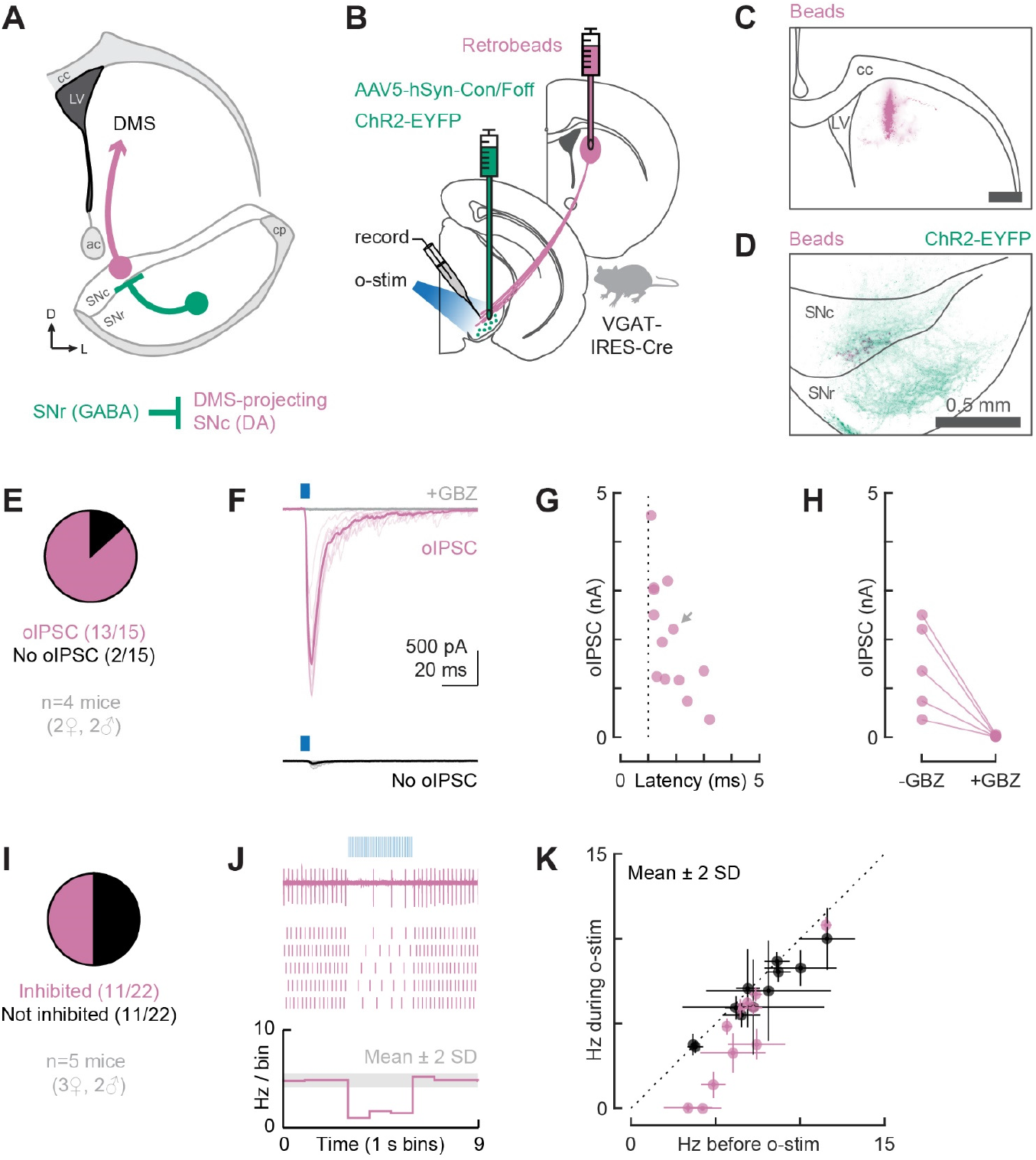
VGAT+ cells in SNr monosynaptically inhibit DMS-projecting DA neurons in SNc and suppress their tonic firing. **(A)** Schematic of the tested circuit. **(B)** Experimental design for probing the connection between VGAT+ cells in SNr and DMS-projecting DA neurons in SNc. All injections were done in adult VGAT-IRES-Cre mice. AAV5-hSyn-Con/Foff-ChR2-EYFP was injected into SNr to deliver ChR2 to VGAT+ cells. Foff is irrelevant in this setup. Retrobeads were injected into DMS to label the soma of DMS-projecting neurons for targeted patching in midbrain slices. The o-stim was delivered via the objective (460 nm, ~20 mW/mm^2^). **(C)** Distribution of retrobeads (magenta) in a representative striatum slice. Scale bar: 0.5 mm. **(D)** Distribution of bead-labeled somas (magenta) and ChR2-EYFP-labeled neuropil (green) in a representative midbrain slice. SNc was outlined based on TH immunolabeling. **(E)** Proportion of DMS-projecting neurons recorded in whole-cell mode that did (magenta) or did not (black) respond to the o-stim (5 ms pulse) with an oIPSC. **(F)** Example recording from a cell that did (magenta) or did not (black) show an oIPSC. The oIPSC was absent after GBZ perfusion (gray). Thin lines: individual sweeps. Thick lines: average. **(G)** oIPSC amplitude and onset latency for all responding cells (dotted line = 1 ms). A gray arrow indicates the oIPSC shown in F. **(H)** oIPSC amplitude before and after GBZ perfusion for all tested cells. **(I)** Proportion of DMS-projecting neurons recorded in loose seal mode that did (magenta) or did not (black) have their tonic firing suppressed by the o-stim (5 ms pulses at 20 Hz for 3 s). **(J)** Example loose seal recording from a cell that was inhibited by the o-stim. Top: raw data from a single sweep. Middle: raster plot showing action potentials from 5 sweeps. Bottom: histogram of the average firing rate across all sweeps. The gray shaded area indicates mean±2SD for the baseline firing rate. Note that the firing rate drops below mean-2SD during the o-stim. **(K)** Average firing rate before and during the o-stim for all cells recorded in loose seal mode (inhibited cells: magenta; not inhibited: black). Error bars represent ±2SD. Dotted line: unity.

To allow for optogenetic stimulation of GABAergic neurons in SNr, we injected an adeno-associated virus (AAV) carrying a Cre-dependent channelrhodopsin-2 (ChR2) construct into the SNr of VGAT-IRES-Cre mice. The specific virus used (AAV5-hSyn-Con/Foff-ChR2-EYFP) also contains a feature by which ChR2 expression is turned off by Flp recombinase. In these initial experiments (Figure 1-2), the Flp-dependent feature is irrelevant. However, it was crucial for later experiments and so we decided to use the same virus throughout this study.

We began by examining SNr inputs to DLS-projecting DA neurons (Figure 1A-B). We verified that all retrobead injections were contained within the DLS (Extended Figure 1A-B). As expected from previous findings (Farassat et al., 2019; Haber et al., 2000; Ikemoto, 2007; Lerner et al., 2015), the resulting bead-labeled DLS-projecting DA cells were located in mid to lateral SNc (Figure 1D and Extended Figure 1C-D). We started by evaluating the proportion of DLS-projecting DA neurons that were monosynaptically inhibited by GABAergic SNr cells. Our goal in this experiment was to maximize the detection of inhibitory post-synaptic currents (IPSCs) and minimize false negative results. Therefore, we recorded from bead-labeled cells in whole-cell mode using a high chloride internal solution (E_CI_ = 0 mV) and held the cells at −70 mV. In addition, we used a pharmacological approach to isolate monosynaptic connections (Petreanu et al., 2009) – we added tetrodotoxin (TTX, 1μM) to the bath to block action potentials, and 4-aminopyridine (4-AP, 100μM) to boost the neurotransmitter release probability from ChR2-expressing terminals. To isolate inhibitory synapses, we added NBQX (5μM) and D-AP5 (50μM) to the bath to block AMPA and NMDA receptor currents, respectively. A 5 ms blue light pulse (460 nm, ~20 mW/mm^2^) was delivered to the slice to stimulate ChR2-expressing terminals. Under this configuration, we found that 69% (18/26) of the recorded DLS-projecting neurons were monosynaptically inhibited by GABAergic SNr cells (Figure 1E-F). The amplitude of the optogenetically-evoked IPSCs (oIPSCs) ranged from 0.2 to 3.9 nA (mean±SD: 1.8±1.1 nA) and the onset latencies were within 5 ms (range: 1.4-2.8 ms; mean±SD: 1.7±0.4 ms), consistent with the isolation of monosynaptic connections (Figure 1G). For all tested cells, the oIPSC was blocked by the GABA_A_ receptor antagonist gabazine (GBZ, 10μM; Figure 1H).

These experiments established a robust synaptic connectivity between SNr and DLS-projecting SNc DA neurons, but the measurements were performed under non-physiological conditions (large chloride driving force and high neurotransmitter release probability). Therefore, we additionally wanted to assess whether the observed GABAergic inputs could suppress the tonic firing of DA cells under more physiological conditions. To avoid manipulating intracellular chloride, we recorded from bead-labeled cells in a loose seal configuration. NBQX and D-AP5 were again added to bath, but not TTX and 4-AP. For these experiments, we used a 3 s-long light train consisting of 5 ms pulses delivered at 20 Hz (460 nm, ~20 mW/mm^2^). A cell was considered inhibited if the light train reduced its firing rate by more than 2 standard deviations (SD) from the mean (Figure 1J-K). Suppression of tonic firing was observed in 68% (19/28) of the recorded DLS-projecting cells (Figure 1I). The percentage of cells whose firing was inhibited by SNr inputs closely matched the percentage in which oIPSCs were observed, arguing that the GABAergic connections detected onto DLS-projecting DA neurons are effective at controlling their firing rates.

We next examined SNr inputs to DMS-projecting DA neurons (Figure 2A-B). We verified that all retrobead injections were contained within the DMS (Extended Figure 2A-B) and that, as expected (Lerner et al., 2015), bead-labeled DMS-projecting DA cells were medially located in SNc (Figure 2D and Extended Figure 2C-D). Under recording conditions used to isolate monosynaptic inhibitory connections, we found that 87% (13/15) of the recorded DMS-projecting neurons were monosynaptically inhibited by GABAergic SNr cells (Figure 2E-F). The oIPSC amplitude ranged from 0.4 to 4.5 nA (mean±SD: 2.0±1.2 nA) and the onset latencies were within 5 ms (range: 1.1-3.2 ms; mean±SD: 1.8±0.7 ms; Figure 2G). For all tested cells, the oIPSC was blocked by GBZ (Figure 2H). Under a loose seal configuration, suppression of tonic firing was observed in 50% (11/22) of the recorded DMS-projecting cells (Figure 2I-K). In contrast to our findings for DLS-projecting DA neurons, we found a higher percentage of DMS-projecting cells receiving monosynaptic inputs from SNr (87% vs 69%), but a lower percentage of DMS-projecting cells whose tonic firing was inhibited by SNr (50% vs 68%).

Collectively, these findings suggest that both DLS- and DMS-projecting DA neurons in SNc receive robust inhibition from GABAergic SNr cells. Although we did not assess disinhibition directly, such robust inhibition suggests that a decrease in the tonic firing rate of GABAergic SNr cells would be sufficient to disinhibit DLS- and DMS-projecting DA cells. Moreover, our results hint at a dissociation between optogenetically-defined synaptic connectivity and effective suppression of tonic firing. Asymmetries in the proportion of connected versus effectively inhibited cells may indicate fundamental differences between sub-circuits involving DLS- and DMS-projecting DA neurons.

### Dissection of Polysynaptic Striato-Nigro-Striatal Circuits Using a Transsynaptic Cre Virus and Intersectional Genetics

The previous experiments assessed two nigro-striatal circuits: SNr→SNc→DLS (Figure 1) and SNr→SNc→DMS (Figure 2). We next wanted to layer on to our assessment of these circuits the contributions of striatal inputs to SNr, which would allow either for striatal neurons to control disinhibition of their own dopaminergic input (through closed loops such as DLS→SNr→SNc→DLS) or for one striatal region to regulate dopaminergic transmission in a neighboring region (e.g., DMS→SNr→SNc→DLS) as proposed in the ascending spiral hypothesis (Haber et al., 2000; Yin and Knowlton, 2006). While hypotheses about DA neuron disinhibition through striato-nigro-striatal circuits are often incorporated into theory (e.g., Lüscher et al., 2020), the difficulty of tracing synaptic connectivity through a polysynaptic circuit has impeded their testability. Therefore, hypotheses about the structure and function of these circuits have remained highly speculative. We realized that new anterograde tracing (Zingg et al., 2017, 2020) and combinatorial targeting tools (Fenno et al., 2014) would – for the first time – allow highly-specific tests of the structure and function of striato-nigro-striatal circuits.

To label SNr cells by their striatal inputs, we used scAAV1-hSyn-Cre as a transsynaptic anterograde Cre vector (Zingg et al., 2020). When injected into DLS or DMS, this virus will transduce SPNs at the injection site and the post-synaptic targets of these SPNs throughout the brain. Thus, cells that receive a monosynaptic input from DLS or DMS will also carry Cre. We refer to these anterogradely-labeled cells as “DLS-targeted” and “DMS-targeted,” respectively. Before continuing our electrophysiology experiments, we examined the resulting histology in striatum and substantia nigra (SN) after injection of scAAV1-hSyn-Cre into striatum (Figure 3).

**Figure 3.**
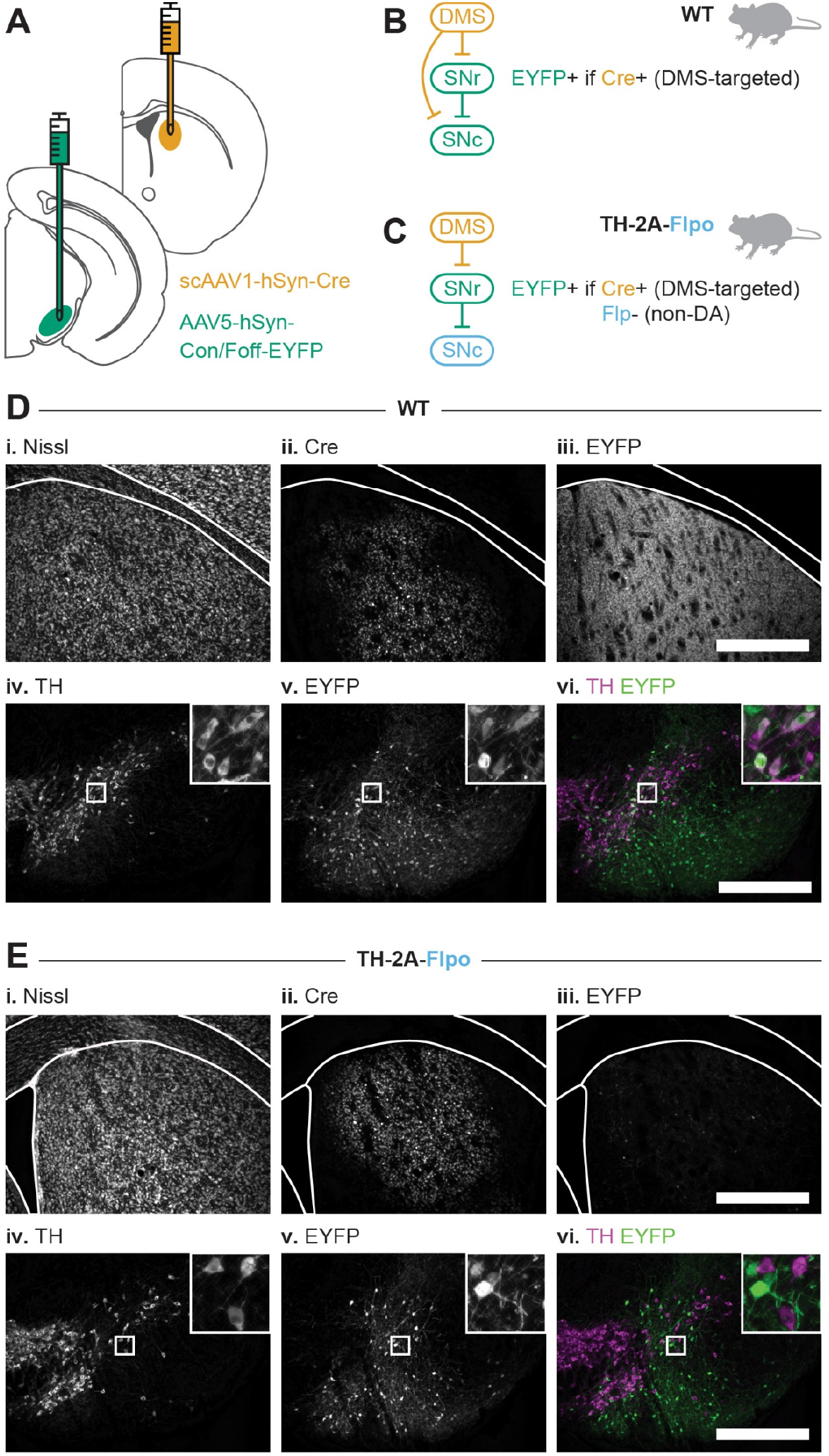
Viral strategy used for polysynaptic circuit dissection. **(A)** Experimental design for labeling DMS-targeted, non-dopaminergic neurons in SNr. scAAV1-hSyn-Cre moves transsynaptically in the anterograde direction and is injected into DMS to deliver Cre to DMS-targeted neurons. AAV5-hSyn-Con/Foff-EYFP is injected into SNr to deliver EYFP to cells that are both Cre+ and Flp-. **(B)** Resulting EYFP labeling in a WT mouse (all cells are Flp-). DMS-targeted cells in SNr and SNc express Cre. Any Cre+ cell carrying Con/Foff-EYFP will express EYFP, regardless of GABAergic or dopaminergic phenotype. **(C)** Resulting EYFP labeling in a TH-2A-Flpo mouse. Dopaminergic cells are Flp+ and will not express EYFP. Consequently, only DMS-targeted, non-dopaminergic cells in SNr express EYFP. **(D-E)** Example histology from the striatum (top row) and SN (bottom row) after injections in WT (D) and TH-2A-Flpo (E) mice. Scale bar: 0.5 mm.

First, we injected wildtype (WT) mice (Figure 3B). We verified that scAAV1-hSyn-Cre did not lesion the striatum, as evidenced by healthy Nissl staining (Figure 3D-i), and observed that Cre expression was restricted to the targeted region (Figure 3D-ii). Next, we looked for Cre expression in SNr. To do so, we injected a Cre-dependent EYFP construct (AAV5-hSyn-Con/Foff-EYFP) into SNr. EYFP+ cells were observed in SNr (Figure 3D-v), but EYFP+ fibers were also observed in the striatum (Figure 3D-iii). GABAergic SNr cells receive monosynaptic inputs from striatum but do not project directly to striatum, whereas dopaminergic SNc neurons do both (Evans et al., 2020; Lerner et al., 2015; Matsuda et al., 2009; Watabe-Uchida et al., 2012; Zingg et al., 2020). Thus, EYFP+ fibers observed within the striatum indicate that DA neurons received Cre. Indeed, after immunostaining for the DA marker tyrosine hydroxylase (TH), we confirmed that EYFP-labeled cells in SN included both TH- and TH+ cells (Figure 3D-vi). It is also possible that some DA neurons received Cre through unintended retrograde movement of the transsynaptic Cre virus (Ii et al., 2008; Zingg et al., 2017, 2020). However, any retrograde movement of the virus does not affect the labeling of SNr neurons, since these cells do not project to striatum (McElvain et al., 2021a).

While not surprising, the finding that SNc DA neurons were labeled with Cre by injection of scAAV1-hSyn-Cre in the striatum presented a problem for our experimental design, which required that we limit ChR2 expression to GABAergic SNr neurons. Therefore, we used an intersectional Cre/Flp recombinase expression strategy to exclude expression of EYFP/ChR2 from DA neurons. Namely, we injected scAAV1-hSyn-Cre into the DMS of TH-2A-Flpo mice, which express Flp recombinase in DA neurons (Poulin et al., 2018). We then injected the same EYFP virus as above (AAV5-hSyn-Con/Foff-EYFP) into SNr. The Con/Foff construct allows expression of EYFP in cells that express Cre, but not Flp. Therefore, we could positively label non-dopaminergic SNr neurons identified as receiving input from a particular striatal subregion (Figure 3C). Using this strategy, we did not find evidence of overlapping EYFP and TH expression in SN (Figure 3E-vi). In addition, we did not observe EYFP+ fibers in the striatum (Figure 3E-iii). The success of this strategy is more easily visualized with an EYFP virus, which labels the cytoplasm of neurons, but this strategy was equally successful when we used a ChR2 virus (AAV5-hSyn-Con/Foff-ChR2-EYFP, Extended Figure 3A).

In sum, we can deliver ChR2 to DMS- and DLS-targeted non-dopaminergic cells in SN with two viral injections in a TH-2A-Flpo mouse: a transsynaptic anterograde Cre virus in striatum (DMS or DLS) and a Con/Foff-ChR2 virus in SNr.

### Characterization of Closed Striato-Nigro-Striatal Loops

By combining retrobead injections in striatum with our viral strategy in TH-2A-Flpo mice, we could investigate the structure and function of multiple striato-nigro-striatal circuits. Because basal ganglia circuits are thought to operate primarily in parallel closed loops (Alexander et al., 1986; Haber et al., 2000; Lee et al., 2020; Mandelbaum et al., 2019; Yin and Knowlton, 2006), we began by testing closed striato-nigro-striatal loops through which DLS and DMS could regulate their own dopaminergic drive.

To test a closed DLS loop (Figure 4A), we injected both the transsynaptic Cre virus (scAAV1-hSyn-Cre) and red retrobeads into the DLS of TH-2A-Flpo mice. We also injected AAV5-hSyn-Con/Foff-ChR2-EYFP into SNr. With this design, we could record from bead-labeled DLS-projecting DA neurons in SNc while optogenetically stimulating DLS-targeted GABAergic neurons in SNr (Figure 4B). We verified that all DLS injections were contained within the DLS (Figure 4C and Extended Figure 4A-B). We also observed that both bead-labeled somas and ChR2-EYFP+ neuropil were located in mid-lateral SN (Figure 4D). Under recording conditions used to isolate monosynaptic inhibitory connections, we found that 53% (9/17) of the recorded DLS-projecting neurons were monosynaptically inhibited by DLS-targeted GABAergic cells in SNr (Figure 4E-F). The oIPSC amplitude ranged from 0.2 to 3.8 nA (mean±SD: 1.6±1.3 nA) and the onset latencies were 1.2-2.8 ms (mean±SD: 1.7±0.5 ms; Figure 4G). For all tested cells, the oIPSC was blocked by GBZ (Figure 4H). Under a loose seal configuration, suppression of tonic firing was observed in 50% (9/18) of the recorded DLS-projecting cells (Figure 4I-K). The percentage of cells whose firing was suppressed closely matched the percentage in which oIPSCs were observed (50% vs 53%), recapitulating the correlation between effective inhibition and synaptic connectivity observed for DLS-projecting DA neurons previously (Figure 1, 68% vs 69%).

**Figure 4.**
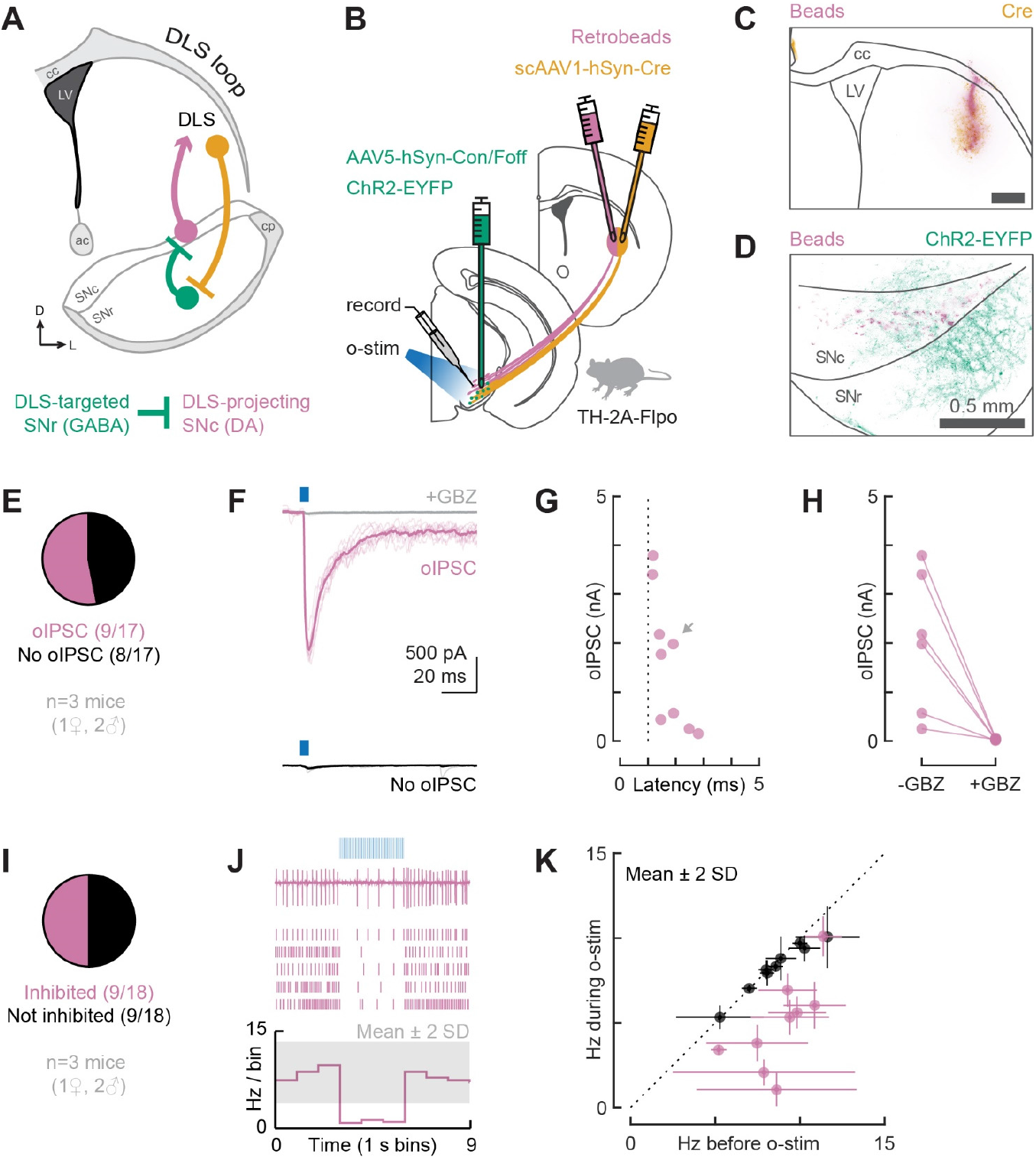
DLS-targeted GABAergic cells in SNr monosynaptically inhibit DLS-projecting DA neurons in SNc and suppress their tonic firing. **(A)** Schematic of the tested circuit. **(B)** Experimental design for probing the connection between DLS-targeted GABAergic cells in SNr and DLS-projecting DA neurons in SNc. All injections were done in adult TH-2A-Flpo mice. The anterograde transsynaptic virus scAAV1-hSyn-Cre was injected into DLS to label DLS-targeted cells with Cre. AAV5-hSyn-Con/Foff-ChR2-EYFP was injected into SNr to deliver ChR2 to cells carrying Cre but not Flp. Retrobeads were injected into DLS to label the soma of DLS-projecting neurons for targeted patching in midbrain slices. The o-stim was delivered via the objective (460 nm, ~20 mW/mm^2^). **(C)** Distribution of retrobeads (magenta) and Cre (yellow) in a representative striatum slice. Scale bar: 0.5 mm. **(D)** Distribution of bead-labeled somas (magenta) and ChR2-EYFP-labeled neuropil (green) in a representative midbrain slice. SNc was outlined based on TH immunolabeling. **(E)** Proportion of DLS-projecting neurons recorded in whole-cell mode that did (magenta) or did not (black) respond to the o-stim (5 ms pulse) with an oIPSC. **(F)** Example recording from a cell that did (magenta) or did not (black) show an oIPSC. The oIPSC was absent after GBZ perfusion (gray). Thin lines: individual sweeps. Thick lines: average. **(G)** oIPSC amplitude and onset latency for all responding cells (dotted line = 1 ms). A gray arrow indicates the oIPSC shown in F. **(H)** oIPSC amplitude before and after GBZ perfusion for all tested cells. **(I)** Proportion of DLS-projecting neurons recorded in loose seal mode that did (magenta) or did not (black) have their tonic firing suppressed by the o-stim (5 ms pulses at 20 Hz for 3 s). **(J)** Example loose seal recording from a cell that was inhibited by the o-stim. Top: raw data from a single sweep. Middle: raster plot showing action potentials from 5 sweeps. Bottom: histogram of the average firing rate across all sweeps. The gray shaded area indicates mean±2SD for the baseline firing rate. Note that the firing rate drops below mean-2SD during the o-stim. **(K)** Average firing rate before and during the o-stim for all cells recorded in loose seal mode (inhibited cells: magenta; not inhibited: black). Error bars represent ±2SD. Dotted line: unity.

We next examined a closed DMS loop (Figure 5A). To do so, we injected both the transsynaptic Cre virus (scAAV1-hSyn-Cre) and red retrobeads into the DMS of TH-2A-Flpo mice. We also injected AAV5-hSyn-Con/Foff-ChR2-EYFP into SNr. This design enabled us to record from bead-labeled DMS-projecting DA neurons in SNc while optogenetically stimulating DMS-targeted GABAergic neurons in SNr (Figure 5B). We verified that all DMS injections were contained within the DMS (Figure 5C and Extended Figure 5A-B). We also observed that both bead-labeled somas and ChR2-EYFP+ neuropil were medially located in SN (Figure 5D). Under recording conditions used to isolate monosynaptic inhibitory connections, we found that 67% (16/24) of the recorded DMS-projecting neurons were monosynaptically inhibited by DMS-targeted GABAergic cells in SNr (Figure 5E-F). The oIPSC amplitude ranged from 0.2 to 3.5 nA (mean±SD: 1.8±1.2 nA) and the onset latencies were 1.1-2.4 ms (mean±SD: 1.6±0.4 ms; Figure 5G). For all tested cells, the oIPSC was blocked by GBZ (Figure 5H). Under a loose seal configuration, suppression of tonic firing was observed in 35% (9/26) of the recorded DMS-projecting cells (Figure 5I-K). The percentage of cells whose firing was inhibited was approximately half of the percentage in which oIPSCs were observed (35% versus 67%), corroborating the dissociation between effective inhibition and synaptic connectivity observed for DMS-projecting DA neurons previously (Figure 2, 50% vs 87%).

**Figure 5.**
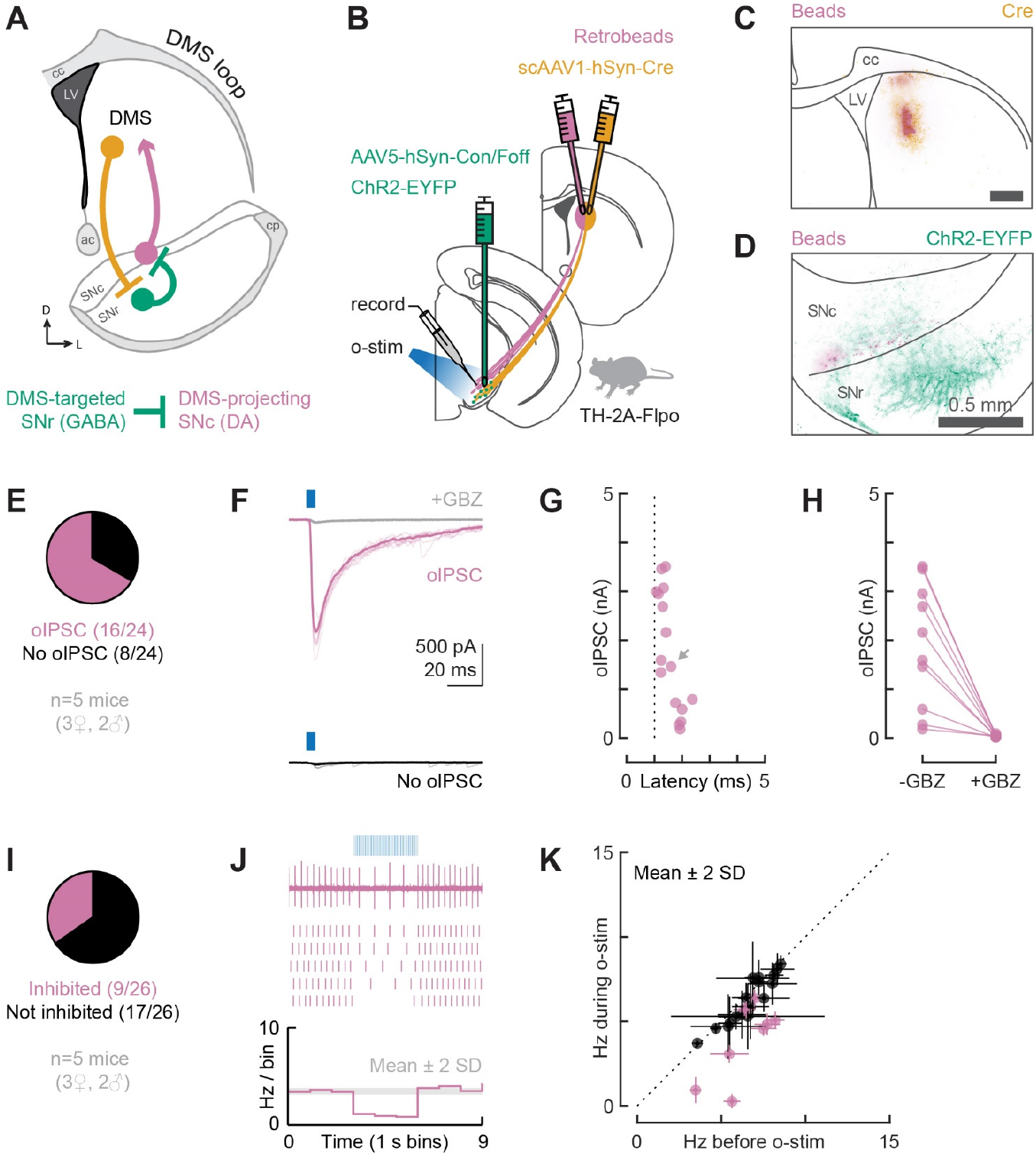
DMS-targeted GABAergic cells in SNr monosynaptically inhibit DMS-projecting DA neurons in SNc and suppress their tonic firing. **(A)** Schematic of the tested circuit. **(B)** Experimental design for probing the connection between DMS-targeted GABAergic cells in SNr and DMS-projecting DA neurons in SNc. All injections were done in adult TH-2A-Flpo mice. The anterograde transsynaptic virus scAAV1-hSyn-Cre was injected into DMS to label DMS-targeted cells with Cre. AAV5-hSyn-Con/Foff-ChR2-EYFP was injected into SNr to deliver ChR2 to cells carrying Cre but not Flp. Retrobeads were injected into DMS to label the soma of DMS-projecting neurons for targeted patching in midbrain slices. The o-stim was delivered via the objective (460 nm, ~20 mW/mm^2^). **(C)** Distribution of retrobeads (magenta) and Cre (yellow) in a representative striatum slice. Scale bar: 0.5 mm. **(D)** Distribution of bead-labeled somas (magenta) and ChR2-EYFP-labeled neuropil (green) in a representative midbrain slice. SNc was outlined based on TH immunolabeling. **(E)** Proportion of DMS-projecting neurons recorded in whole-cell mode that did (magenta) or did not (black) respond to the o-stim (5 ms pulse) with an oIPSC. **(F)** Example recording from a cell that did (magenta) or did not (black) show an oIPSC. The oIPSC was absent after GBZ perfusion (gray). Thin lines: individual sweeps. Thick lines: average. **(G)** oIPSC amplitude and onset latency for all responding cells (dotted line = 1 ms). A gray arrow indicates the oIPSC shown in F. **(H)** oIPSC amplitude before and after GBZ perfusion for all tested cells. **(I)** Proportion of DMS-projecting neurons recorded in loose seal mode that did (magenta) or did not (black) have their tonic firing suppressed by the o-stim (5 ms pulses at 20 Hz for 3 s). **(J)** Example loose seal recording from a cell that was inhibited by the o-stim. Top: raw data from a single sweep. Middle: raster plot showing action potentials from 5 sweeps. Bottom: histogram of the average firing rate across all sweeps. The gray shaded area indicates mean±2SD for the baseline firing rate. Note that the firing rate drops below mean-2SD during the o-stim. **(K)** Average firing rate before and during the o-stim for all cells recorded in loose seal mode (inhibited cells: magenta; not inhibited: black). Error bars represent ±2SD. Dotted line: unity.

Collectively, these findings confirm the existence of closed striato-nigro-striatal loops through which DLS and DMS could alter their own dopaminergic drive via inhibition of GABAergic cells in SNr and disinhibition of DA cells in SNc. Moreover, our findings suggest that DLS would be more effective at such disinhibition than DMS.

### Open Spiral Striato-Nigro-Striatal Circuits Are Unlikely to Support Robust Dopamine Disinhibition

After employing our experimental strategy to test closed striato-nigro-striatal loops, we used a similar approach to test open-loop spiral circuits, beginning with the ascending spiral circuit (DMS→SNr→SNc→DLS). We injected the transsynaptic Cre virus (scAAV1-hSyn-Cre) into the DMS and red retrobeads into the DLS of TH-2A-Flpo mice (Figure 6A). We also injected AAV5-hSyn-Con/Foff-ChR2-EYFP into SNr. With this design, we could record from bead-labeled DLS-projecting DA neurons in SNc while optogenetically stimulating DMS-targeted GABAergic neurons in SNr (Figure 6B). We verified that all injections in striatum were contained within their target areas (Figure 6C and Extended Figure 6A-B). We also observed an overlap of bead-labeled cells and ChR2-EYFP+ neuropil in SN (Figure 6D), consistent with the predictions of the ascending spiral hypothesis (Haber et al., 2000). Under recording conditions used to isolate monosynaptic inhibitory connections, we found that 50% (15/30) of the recorded DLS-projecting neurons were monosynaptically inhibited by DMS-targeted GABAergic cells in SNr (Figure 5E-F). The oIPSC amplitude ranged from 0.1 to 3.3 nA (mean±SD: 0.9±0.9 nA) and the onset latencies were 1.1–4.3 ms (mean±SD: 1.8±0.8 ms; Figure 6G). For all tested cells, the oIPSC was blocked by GBZ (Figure 6H). Under a loose seal configuration, however, suppression of tonic firing was NOT observed in any of the recorded DLS-projecting cells (0/23, Figure 6I-K). The striking mismatch between the percentage of cells whose firing was inhibited and the percentage in which oIPSCs were observed was unexpected and in stark contrast to the nearly perfect match between synaptic connectivity and effective inhibition for DLS-projecting cells in our previous experiments (Figure 1 and Figure 4). Our findings suggest that there is a fundamental difference between the closed DLS loop and the ascending spiral connecting DMS to DLS. Although synaptic connections exist at roughly similar rates in the two circuits (53% vs 50%), the ability of these circuits to control the tonic firing of DA neurons is remarkably different (50% vs 0%).

**Figure 6.**
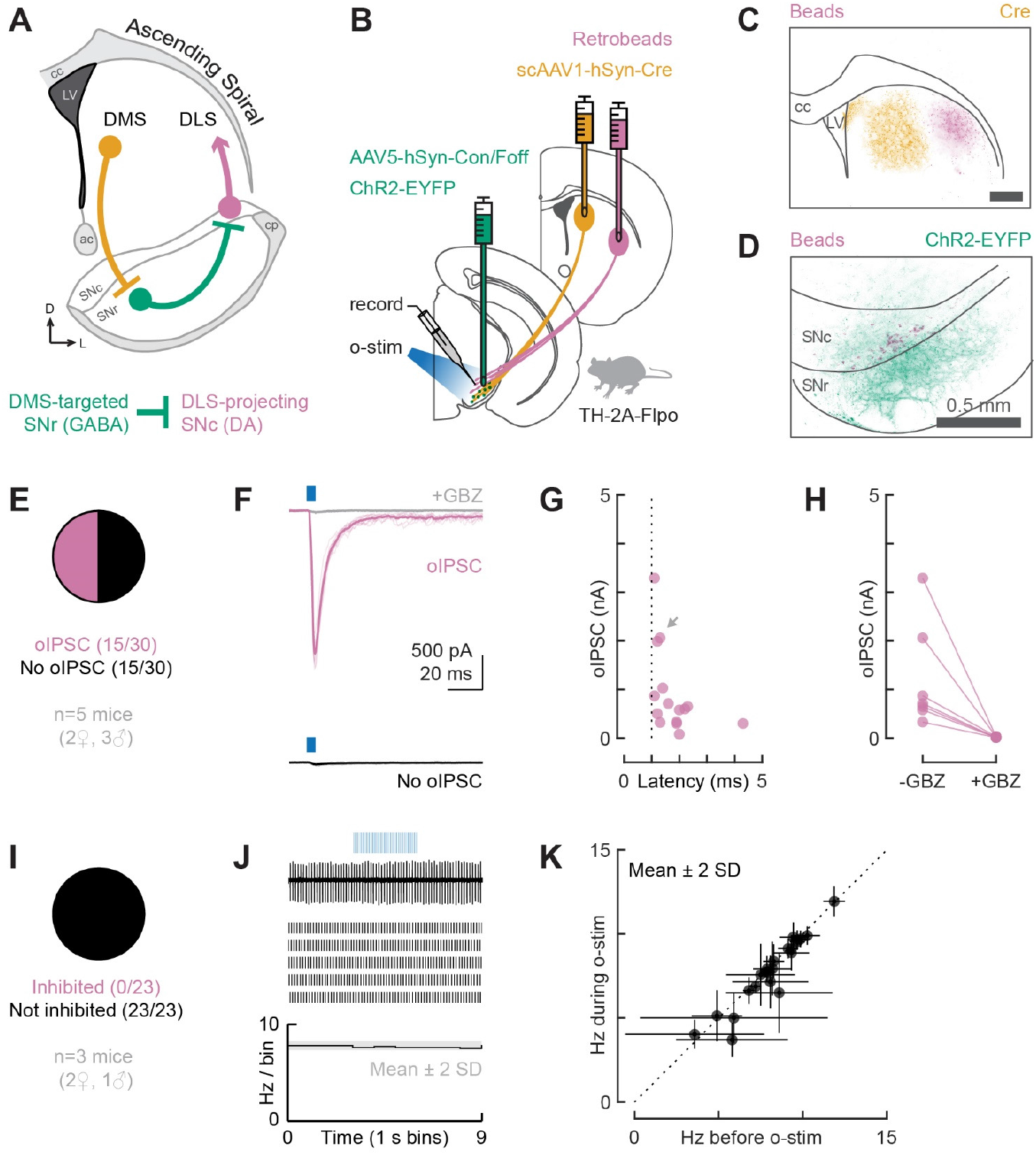
DMS-targeted GABAergic cells in SNr monosynaptically inhibit DLS-projecting DA neurons in SNc but do not suppress their tonic firing. **(A)** Schematic of the tested circuit. **(B)** Experimental design for probing the connection between DMS-targeted GABAergic cells in SNr and DLS-projecting DA neurons in SNc. All injections were done in adult TH-2A-Flpo mice. The anterograde transsynaptic virus scAAV1-hSyn-Cre was injected into DMS to label DMS-targeted cells with Cre. AAV5-hSyn-Con/Foff-ChR2-EYFP was injected into SNr to deliver ChR2 to cells carrying Cre but not Flp. Retrobeads were injected into DLS to label the soma of DLS-projecting neurons for targeted patching in midbrain slices. The o-stim was delivered via the objective (460 nm, ~20 mW/mm^2^). **(C)** Distribution of retrobeads (magenta) and Cre (yellow) in a representative striatum slice. Scale bar: 0.5 mm. **(D)** Distribution of bead-labeled somas (magenta) and ChR2-EYFP-labeled neuropil (green) in a representative midbrain slice. SNc was outlined based on TH immunolabeling. **(E)** Proportion of DLS-projecting neurons recorded in whole-cell mode that did (magenta) or did not (black) respond to the o-stim (5 ms pulse) with an oIPSC. **(F)** Example recording from a cell that did (magenta) or did not (black) show an oIPSC. The oIPSC was absent after GBZ perfusion (gray). Thin lines: individual sweeps. Thick lines: average. **(G)** oIPSC amplitude and onset latency for all responding cells (dotted line = 1 ms). A gray arrow indicates the oIPSC shown in F. **(H)** oIPSC amplitude before and after GBZ perfusion for all tested cells. **(I)** Proportion of DLS-projecting neurons recorded in loose seal mode that did (magenta) or did not (black) have their tonic firing suppressed by the o-stim (5 ms pulses at 20 Hz for 3 s). **(J)** Example loose seal recording from a cell that was not inhibited by the o-stim. Top: raw data from a single sweep. Middle: raster plot showing action potentials from 5 sweeps. Bottom: histogram of the average firing rate across all sweeps. The gray shaded area indicates mean±2SD for the baseline firing rate. Note that the firing rate stays above mean-2SD during the o-stim. **(K)** Average firing rate before and during the o-stim for all recorded cells (inhibited cells: magenta; not inhibited: black). Error bars represent ±2SD. Dotted line: unity.

In previous work establishing the ascending spiral hypothesis, a lack of overlap between axons from lateral striatum and the cell bodies of SN neurons projecting to medial striatum was noted (Haber et al., 2000). This result led to the prediction that there is limited connectivity in a “descending” spiral (DLS→SNr→SNc→DMS), yet this prediction has not been tested. Indeed, such overlap is not necessary for the existence of a functional polysynaptic circuit. To examine the descending spiral circuit (Figure 7A), we injected the transsynaptic Cre virus (scAAV1-hSyn-Cre) into the DLS and red retrobeads into the DMS of TH-2A-Flpo mice. We also injected AAV5-hSyn-Con/Foff-ChR2-EYFP into SNr. With this design, we could record from bead-labeled DMS-projecting DA neurons in SNc while optogenetically stimulating DLS-targeted GABAergic neurons in SNr (Figure 7B). We verified that all injections in striatum were contained within their target areas (Figure 7C and Extended Figure 7A-B). We also observed poor overlap of bead-labeled cells and ChR2-EYFP+ neuropil in SN (Figure 7D), again consistent with the predictions of the ascending spiral hypothesis (Haber et al., 2000). However, despite the lack of overlap, we found that 45% (13/29) of the recorded DMS-projecting neurons were monosynaptically inhibited by DLS-targeted GABAergic cells in SNr (Figure 7E-F). The oIPSC amplitude ranged from 0.2 to 3.6 nA (mean±SD: 1.5±1.1 nA) and the onset latencies were 1.2–4.6 ms (mean±SD: 2.1±1.1 ms; Figure 7G). For all tested cells, the oIPSC was blocked by GBZ (Figure 7H). The connectivity we observed was surprising. However, we did not observe much inhibition of tonic firing through these synaptic connections. Under a loose seal configuration, suppression of tonic firing was observed in only 4% (1/26) of the recorded DMS-projecting cells (Figure 7I-K). The striking mismatch between synaptic connectivity and inhibition of tonic firing was once again unexpected, but not as surprising, given that some mismatch was previously observed for DMS-projecting cells (Figure 2 and Figure 5). Together, our results from testing the ascending and descending spiral circuits suggest that these circuits are unlikely to support robust DA neuron disinhibition, at least in naïve mice.

**Figure 7.**
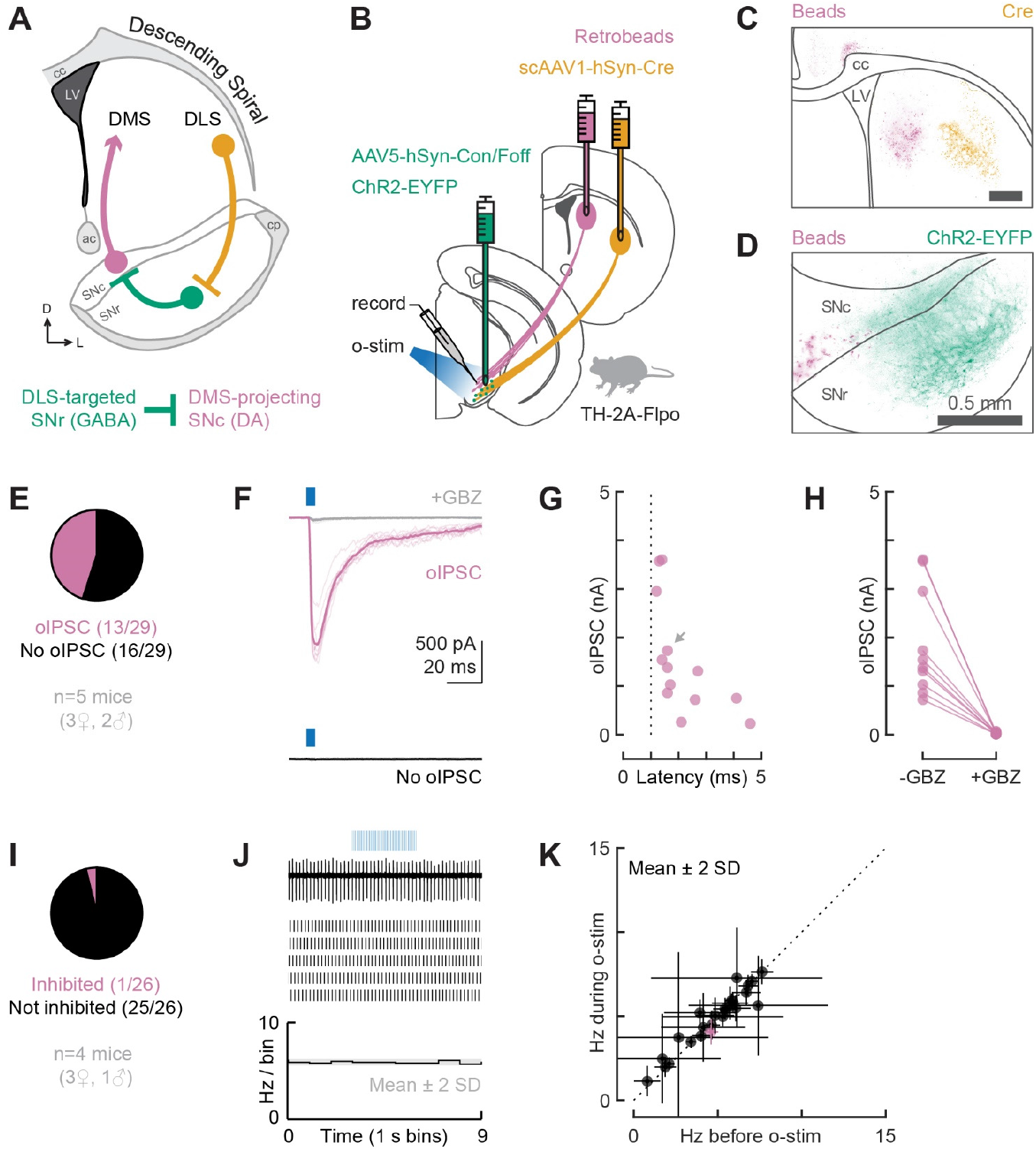
DLS-targeted GABAergic cells in SNr monosynaptically inhibit DMS-projecting DA neurons in SNc but do not suppress their tonic firing. **(A)** Schematic of the tested circuit. **(B)** Experimental design for probing the connection between DLS-targeted GABA cells in SNr and DMS-projecting DA neurons in SNc. All injections were done in adult TH-2A-Flpo mice. The anterograde transsynaptic virus scAAV1-hSyn-Cre was injected into DLS to label DLS-targeted cells with Cre. AAV5-hSyn-Con/Foff-ChR2-EYFP was injected into SNr to deliver ChR2 to cells carrying Cre but not Flp. Retrobeads were injected into DMS to label the soma of DMS-projecting neurons for targeted patching in midbrain slices. The o-stim was delivered via the objective (460 nm, ~20 mW/mm^2^). **(C)** Distribution of retrobeads (magenta) and Cre (yellow) in a representative striatum slice. Scale bar: 0.5 mm. **(D)** Distribution of bead-labeled somas (magenta) and ChR2-EYFP-labeled neuropil (green) in a representative midbrain slice. SNc was outlined based on TH immunolabeling. **(E)** Proportion of DMS-projecting neurons recorded in whole-cell mode that did (magenta) or did not (black) respond to the o-stim (5 ms pulse) with an oIPSC. **(F)** Example recording from a cell that did (magenta) or did not (black) show an oIPSC. The oIPSC was absent after GBZ perfusion (gray). Thin lines: individual sweeps. Thick lines: average. **(G)** oIPSC amplitude and onset latency for all responding cells (dotted line = 1 ms). A gray arrow indicates the oIPSC shown in F. **(H)** oIPSC amplitude before and after GBZ perfusion for all tested cells. **(I)** Proportion of DMS-projecting neurons recorded in loose seal mode that did (magenta) or did not (black) have their tonic firing suppressed by the o-stim (5 ms pulses at 20 Hz for 3 s). **(J)** Example loose seal recording from a cell that was not inhibited by the o-stim. Top: raw data from a single sweep. Middle: raster plot showing action potentials from 5 sweeps. Bottom: histogram of the average firing rate across all sweeps. The gray shaded area indicates mean±2SD for the baseline firing rate. Note that the firing rate stays above mean-2SD during the o-stim. **(K)** Average firing rate before and during the o-stim for all cells recorded in loose seal mode (inhibited cells: magenta; not inhibited: black). Error bars represent ±2SD. Dotted line: unity.

### Strong GABAergic SNr Inputs Onto DA Neurons Do Not Predict Inhibition of Tonic Firing

In both open- and closed-loop striato-nigro-striatal circuits, we observed robust GABAergic connectivity from SNr neurons onto DA SNc neurons, mediated by GABA_A_ receptor transmission. Given this connectivity, and the fact that the amplitude of the recorded oIPSCs was similar in all circuit configurations (Extended Figure 8), we expected to observe similar rates of suppression of DA neuron firing across conditions. Surprisingly, we found instead that the tonic firing of DA neurons was clearly inhibited in closed loops but not open spirals (Figure 8A). Collectively, these findings support a model in which only closed striato-nigro-striatal loops support strong firing suppression, although latent functional connectivity is present in open spirals (Figure 8B). The mechanisms underlying our observed dissociations between monosynaptic connectivity and firing suppression will be an important subject of future studies, which may allow us to explain the conditions under which striatal subregions do and do not communicate with each other via DA circuits.

**Figure 8.**
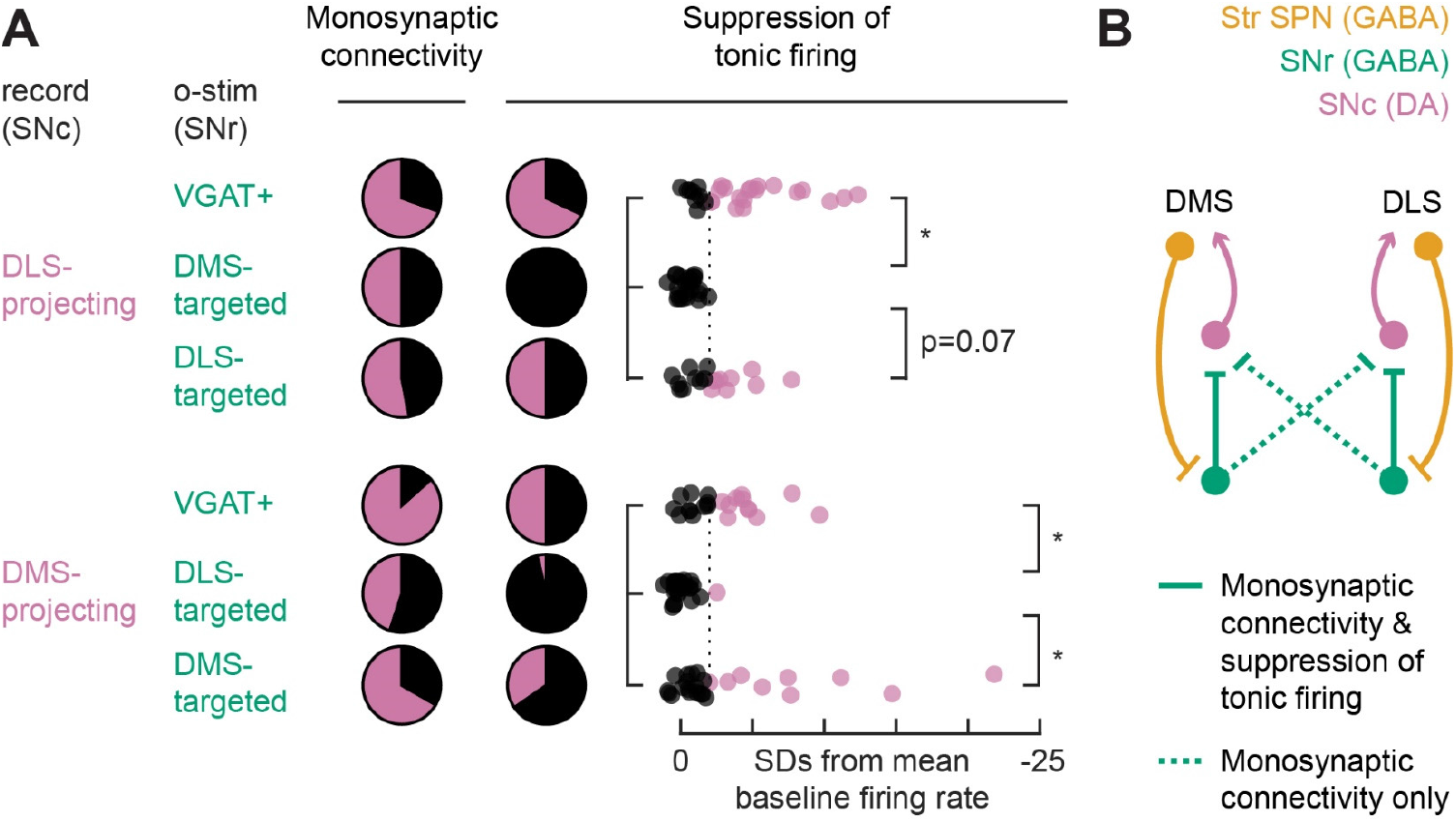
Ascending and descending spirals connecting DMS and DLS are supported by monosynaptic connectivity data but not by suppression of tonic firing. **(A)** Left: monosynaptic connectivity data as proportion of bead-labeled neurons that did (magenta) or did not (black) respond to the o-stim with an oIPSC (reproduced from Figures 1-2E and 4-7E). Middle: proportion of bead-labeled neurons that did (magenta) or did not (black) have their tonic firing suppressed by the o-stim (reproduced from Figures 1-2I and 4-7I). Right: optogenetically-evoked change in tonic firing of bead-labeled cells expressed as SDs from baseline mean (adapted from Figures 1-2K and 4-7K). The dotted line indicates −2SD, our threshold for labeling cells as inhibited (magenta) or not (black) by the o-stim. *p<0.01 (Kruskal-Wallis test followed by Multiple Comparison Test). **(B)** Circuit diagram supported by the data.

## DISCUSSION

### Evidence For and Against the Ascending Spiral Hypothesis

In this paper, we tested multiple striato-nigro-striatal loops connecting two striatal subregions (DMS and DLS) via SNr and SNc (i.e., DMS/DLS→SNr→SNc→DMS/DLS). These loops have the potential to transform activity in a striatal subregion into DA release in the same or neighboring region of striatum by disinhibiting dopaminergic neurons in SNc. We were particularly interested in testing the predictions of the ascending spiral hypothesis, which argues that open loop striato-nigro-striatal circuits permit the progressive disinhibition of DA neurons in a unidirectional, “ascending” (medio-lateral) direction (Haber et al., 2000). We focused on circuits involving the dorsal striatum given that the ascending spiral hypothesis is frequently invoked to explain changes in DMS and DLS that occur over the course of extended training, as animals become proficient in motor skill tasks and/or transition from goal-directed to habitual behavior.

To assess connectivity in the polysynaptic striato-nigro-striatal circuits in question, we labeled GABAergic SNr cells based on their inputs and dopaminergic SNc cells based on their outputs. We then assessed the synaptic connectivity between input-labeled SNr cells and output-labeled SNc cells that belong to a variety of striato-nigro-striatal circuits, including closed and open loops. Our data support the existence of a DMS→SNr→SNc→DLS circuit as predicted by the ascending spiral hypothesis, but challenge the prediction that this circuit alone can support disinhibition in DA neurons (Figure 6). Instead, our data suggest that closed striato-nigro-striatal loops (i.e., DMS→SNr→SNc→DMS and DLS→SNr→SNc→DLS) are better suited to support disinhibition (Figures 4-5). These data for dorsal striatal circuits are complemented by other findings in ventral striatal circuits, which also suggest that disinhibition operates primarily in closed loops between striatal subregions and the dopaminergic midbrain (Yang et al., 2018). Our findings further diverge from the ascending spiral hypothesis by confirming the existence of a descending spiral (DLS→SNr→SNc→DMS) of approximately equal strength to the ascending spiral, challenging the claim of unidirectional information flow (Figure 7).

These results are important because previous anatomical findings about the topography of striato-nigro-striatal circuits (Haber et al., 2000) have inspired the field to interpret behavioral and neural activity findings under the framework of an ascending spiral (Lerner, 2020; Lüscher et al., 2020; Yin and Knowlton, 2006). Indeed, the sequential recruitment of DMS and DLS during motor skill learning and habit formation fits nicely with the ascending spiral hypothesis (Gremel and Costa, 2013; Thorn et al., 2010; Yin et al., 2009). So does the dependence of habit formation on DA projections to DLS (Faure et al., 2005) and the increasing recruitment of DLS DA activity with drug use (Belin and Everitt, 2008; Willuhn et al., 2012). In addition, modeling studies point to striato-nigral circuits in the form of Str→SNr→SNc as a robust means of disinhibition and burst firing in dopaminergic neurons (Lobb et al., 2011). Although no direct evidence exists for impaired DA release in DLS following DMS lesions, ventromedial striatum (VMS) lesions are reported to impair DLS DA release (Willuhn et al., 2012).

While the above findings are consistent with the ascending spiral hypothesis, direct evidence for a continuous polysynaptic circuit connecting DMS→SNr→SNc→DLS was lacking, and other findings do not fit. For instance, if repeated activation of DMS is required to elicit DA release in DLS and drive motor learning and habit formation, one would expect DMS lesions to hinder these processes, but that is not the case. Instead, DMS lesions do not prevent motor skill learning (Yin et al., 2009) and are in fact reported to accelerate habit formation (Gremel and Costa, 2013; Yin et al., 2004, 2005a, 2005b). It is also hard to reconcile the slow time course of habit formation and the associated changes in DLS (days to weeks) with a tri-synaptic circuit theoretically capable of regulating DA release in DLS within tens of milliseconds. One possible explanation is that a disinhibitory ascending spiral circuit is not fully functional in naïve animals but develops slowly during training. The synaptic connections we observed in the DMS→SNr→SNc→DLS circuit could undergo plasticity and/or regulate the plasticity of other inputs onto DA neurons over the course of training even if they do not regulate DA neuron firing in naïve mice.

### Technical Considerations

Two technical caveats could result in underestimation of the connectivity probabilities reported here: (1) incomplete penetrance of our labeling methods and (2) severing of the distal dendrites of DA neurons in midbrain slices. Although our labeling methods are not 100% penetrant, any underestimation due to this caveat should affect all tested circuits similarly since we used the same viruses and retrobeads in all experiments. The severing of distal dendrites, on the other hand, could disproportionately affect some circuit configurations. The substantia nigra has a complex 3D structure that is not fully preserved in coronal slices (Gerfen et al., 1987; Maurin et al., 1999), and slicing could sever the distal dendrites of DA neurons. If a particular sub-population of SNr cells targets these distal dendrites, then this connection is more likely to be underestimated. Alternatively, if DA neurons projecting to DLS or DMS belong predominantly to ventral tier SNc and have a prominent distal dendrite in SNr (Gerfen et al., 1987), connections onto these cells are also more likely to be underestimated. Fortunately, these caveats do not seem to significantly bias our results, given that oIPSCs of similar amplitudes were detected in all circuit configurations (Extended Figure 8). In addition, we assessed synaptic connectivity and effects on tonic firing in slices from the same mice. DA cells recorded in a loose seal configuration that were not inhibited by the optogenetic stimulation (o-stim) were often neighboring DA cells recorded in whole-cell mode that exhibited robust oIPSCs. Hence, the dissociation between synaptic connectivity and effective inhibition reported here is not due to variability in slicing and/or ChR2 expression across animals. We were also careful to sample bead-labeled cells across the entire volume of SN to avoid any biases regarding the medio-lateral, rostro-caudal, or dorsal-ventral location of DA neurons (Extended Figures 1-2 and 4-7). We did not observe any clear correlations between cell location and likelihood of connection for any of the tested circuits.

### Alternatives to the Ascending Spiral Hypothesis

The ascending spiral hypothesis as formulated here is not the only means by which striatal subregions could influence each other. For example, VMS can modulate DLS activity via a long polysynaptic loop through SNr, thalamus and motor cortex (Aoki et al., 2019), bypassing not only DMS but also DA neurons. Other mechanisms might exist through lateral inhibition amongst SPNs (Burke et al., 2017), striatal interneuron networks (Cai and Ford, 2018; Dorst et al., 2020; Fino et al., 2018; Holly et al., 2019; Xu et al., 2015), modulation of DA axon terminals (Kramer et al., 2020; Liu et al., 2021; Mohebi et al., 2019), or through striatal astrocyte networks (Khakh, 2019). Thus, even if the ascending spiral circuit for DMS-DLS communication through the control of DA neuron activity is weak, other circuits may instead support information transfer between DMS and DLS.

### A Dissociation Between Connectivity and Firing Rate Modulation

A key assumption in systems neuroscience is that synaptic connectivity can be used to reverse engineer neural circuit function and dynamics, but our data highlights a dissociation between connectivity and effects on tonic firing. This was surprising, but not completely unexpected, given that monosynaptic connectivity was assessed under conditions designed to maximize detection sensitivity, but not optimized to make physiological measurements. Indeed, oIPSCs were recorded in DA neurons with an uncharacteristically large driving force for chloride, while our loose seal recordings preserved the physiological driving force. DA neurons, which do not express the chloride extruder KCC2 (potassium-chloride cotransporter 2), have a weakly hyperpolarizing chloride reversal potential (Gulácsi et al., 2003). Therefore, inhibition through GABA_A_ receptor activity is primarily due to shunting inhibition and will be less effective at regulating firing rates if synapses are located far from the action potential generating mechanisms of the DA cell. In other words, one might expect lower rates of firing modulation as compared to rates of monosynaptic connectivity if synapses are located on distal dendrites. However, what was most unexpected was that the dissociation between connectivity and firing was not the same in all striato-nigro-striatal circuits tested. A dissociation was found in all striato-nigro-striatal circuits involving DMS-projecting DA neurons, but only one configuration involving DLS-projecting DA neurons: the ascending spiral circuit (Figure 8). Therefore, our hypothesis is that SNr GABAergic synapses participating in the ascending spiral are arranged in fundamentally different patterns from SNr GABAergic synapses NOT participating in the ascending spiral.

Compartmentalization of synaptic inputs has been previously reported for midbrain DA neurons, as has heterogeneity in intrinsic properties (Evans et al., 2017, 2020; Farassat et al., 2019; Lammel et al., 2008, 2011; Tarfa et al., 2017). Notably, striosome SPNs from dorsal striatum target the distal SNr dendrite of SNc DA neurons, while neurons of the globus pallidus external segment (GPe) target the soma and proximal dendrites of DA neurons (Evans et al., 2020). Moreover, striosomes target ventral tier DA neurons, which have a prominent sag current and after-depolarization that support rebound firing (Evans et al., 2017, 2020). Interactions between intrinsic properties and preferential targeting could explain the differences we observed between closed and open loops. Additional layers of synaptic input integration would be possible if, like hippocampal and cortical neurons, DA cells could maintain a compartmentalized responsiveness to GABAergic inputs due to subcellular variance in intracellular chloride (Khirug et al., 2008; Rahmati et al., 2021).

Presynaptic mechanisms could also explain the dissociation we observed. During the detection of oIPSCs, we used 4-AP to boost neurotransmitter release probability from GABAergic SNr cells, but a more physiological release probability was preserved during loose seal recordings. If release probability from GABAergic SNr cells is low, no suppression of tonic firing would be observed in DA neurons even with robust connectivity, unless there were many synapses converging onto the same DA neuron. On the other hand, a high release probability would lead to oIPSC depression during the light train, resulting in brief suppression of tonic firing that is not sustained over the course of seconds. SNr neurons are electrophysiologically heterogeneous (McElvain et al., 2021b) and it is likely that fundamental differences between SNr GABAergic synapses engaged in closed versus open loops stem from the recruitment of non-overlapping subpopulations of GABAergic SNr cells.

### Balancing Striatal Inhibition and Disinhibition of Dopamine Neurons

Given the findings described here regarding the *indirect* connections between striatum and SNc via SNr, and previous research on the *direct* connections between striatum and SNc, it is hard to predict which patterns of striatal activity would support the disinhibition of DA neurons *in vivo*. Multiple rabies tracing studies have characterized the monosynaptic inputs onto projection-defined DA neurons and identified the striatum as a major source of direct inhibition to DA cells (Lerner et al., 2015; Menegas et al., 2015; Watabe-Uchida et al., 2012). However, these direct connections were excluded from computational models of striato-nigro-striatal circuits that predicted disinhibition of DA neurons following striatal activation (Lobb et al., 2011). Lerner and colleagues further dissected these *direct* striato-nigro-striatal circuits with slice electrophysiology and found that DMS preferentially targets DMS-projecting DA neurons, while DLS targets both DMS- and DLS-projecting DA neurons. Thus, monosynaptic connections between striatum and SNc support the existence of closed loops (DMS→SNc→DMS and DLS→SNc→DLS), as well as a descending circuit (DLS→SNc→DMS). Work in ventral striatal circuits also draws attention to the role of direct inhibition of DA neurons by striatal inputs, which can be mediated by GABA_B_ as well as GABA_A_ receptors (Yang et al., 2018). Further investigation is required to compare the relative strength of *direct* and *indirect* striato-nigro-striatal circuits on the activity of DA neurons and test the conditions that favor disinhibition over inhibition *in vivo*.

Finally, it is possible that the balance between inhibition and disinhibition of DA neurons is altered by training, either by synaptic plasticity or by the recruitment of additional circuits during learning. Here, we tested naïve animals (as did Haber et al., 2000 in their original tracing study), but it is possible that different connectivity patterns in striato-nigro-striatal circuits would emerge after training. We and others (Hamid et al., 2021; Seiler et al., 2020; Willuhn et al., 2012) have observed that the *in vivo* patterns of DA axon activity and DA release in DMS and DLS change with training. The reason why DA signaling changes with training is not yet clear, but one exciting possibility is that plasticity in either direct or indirect striato-nigro-striatal circuits is responsible. With the approaches developed here, and with additional innovations to adapt them for *in vivo* investigations, we can begin to rigorously address this hypothesis.

## METHODS

### Animals

Male and female C57BL/6J mice were group housed under a conventional 12:12 h light/dark cycle with ad libitum access to food and water. The VGAT-IRES-Cre knock-in strain was obtained from The Jackson Laboratory (JAX#028862) and the TH-2A-Flpo line was a gift from Dr. Awatramani (Poulin et al., 2018). Animals were crossed in house, and only heterozygous transgenic mice were used for experiments. WT mice used in Figure 3 and Extended Figure 3 were Flp- mice from our TH-2A-Flpo breeding. All experiments were approved by the Northwestern University Institutional Animal Care and Use Committee.

### Stereotaxic surgery

Surgery was performed on adult (7-20 weeks old) male and female mice. Briefly, anesthesia was induced and maintained with isoflurane 1-4% (Patterson Scientific Link 7). Buprenorphine SR (0.5 mg/kg, Zoopharm) and Carprofen (5 mg/kg, Zoetis) were administered subcutaneously for analgesia. Ophthalmic ointment (Puralube, Dechra) was used to prevent dehydration of the cornea. A far infrared heating pad (Kent Scientific) was placed on top of the stereotax (Stoelting 51733D) to keep body temperature at ~37°C. Fur was removed with Nair; 10% povidone-iodine and 70% isopropyl alcohol were used to disinfect the scalp. A small (~1 cm) scalp incision was made to expose the skull, which was later closed with non-absorbable sutures (Ethicon, 661H) and tissue adhesive (Vetbond, 3M). Bregma and lambda were used as landmarks to level the head and guide injections. To drill skull holes, a micromotor drill (Stoelting, 51449) was moved to the appropriate coordinates with the aid of a digital stereotaxic display. Viruses and/or retrobeads were injected into the brain at 50-100 nl/min through a blunt 33-gauge needle using a syringe pump (World Precision Instruments). The needle was left in place for 5 min following the end of the injection, then slowly retracted to avoid leakage up the injection tract. The following coordinates were used (AP, ML, DV – in mm): DMS (0.8, 1.5, −2.8), DLS (0.3, 2.5, −3.3), and SNr (−3.3, 1.2, −4.7). Where indicated, we injected 250 nl of scAAV1-hSyn-Cre (2.81e13 vg/ml, WZ Biosciences) into DMS/DLS, and 250 nl of AAV5-hSyn-Con/Foff-EYFP (2.6e12 vg/ml, UNC, Addgene plasmid #55651) or AAV5-hSyn-Con/Foff-hChR2(H134R)-EYFP (5.3e12 vg/ml, UNC, Addgene plasmid #55646) into SNr. Red retrobeads (LumaFluor) were diluted 1:4 (dilution factor) in sterile saline, and 100 nl were injected into DMS/DLS. When retrobeads were mixed with scAAV1-hSyn-Cre for investigation of closed loops, they were diluted 1:8 in a virus aliquot, and a total volume of 250 nl was injected into DMS or DLS. As a consequence, approximately the same amount of beads was injected into striatum (half the concentration at ~double the volume), and the transsynaptic Cre virus was only slightly diluted (7:8 dilution factor). After surgery, animals were placed on a warm recovery bin until ambulant. A moist nutritional supplement (DietGet 31M, Clear H_2_O) was placed on the floor of the homecage to aid recovery from surgery. 4-6 weeks after surgery, animals received a lethal intraperitoneal injection of Euthasol (1 mg/kg, Virbac), a combination of sodium pentobarbital (390 mg/ml) and sodium phenytoin (50 mg/ml), and underwent a transcardial perfusion for electrophysiology and/or histology experiments.

### Electrophysiology

We followed the methods described by Ting and colleagues (Ting et al., 2014) to prepare acute brain slices from adult mice. Following Euthasol injection, unresponsive mice were transcardially perfused with ice-cold N-Methyl-D-Glucamine (NMDG) artificial cerebrospinal fluid (ACSF) containing (in mM): 92 NMDG, 2.5 KCl, 1.2 NaH2PO4, 30 NaHCO3, 20 HEPES, 25 Glucose, 5 Na-Ascorbate, 2 Thiourea, 3 Na-Pyruvate, 10 MgSO4, 0.5 CaCl2 (Millipore Sigma). All extracellular solutions used for electrophysiology were saturated with 95%O_2_/5%CO_2_ and their pH and osmolarity were adjusted to 7.3-7.4 and 300±5 mOsm, respectively. After perfusion, the brain was quickly removed and cut coronally to separate the rostral half (containing striatum) from the caudal half (containing SN). The cut face of each brain half was glued (Loctite 454) to a specimen holder and immersed into ice-cold NMDG ACSF. Coronal slices (300 μm thick) were made using a vibratome (Leica, VT1200S) set to 0.08 mm/s speed and 1.00 mm amplitude. Striatal slices were saved to confirm injection sites, while midbrain slices were used for recordings. Slices were allowed to recover for 45 min in three 15 min baths: (1) warm (33°C) NMDG ACSF; (2) warm (33°C) recovery ACSF, containing (in mM): 92 NaCl, 2.5 KCl, 1.2 NaH2PO4, 30 NaHCO3, 20 HEPES, 25 Glucose, 5 Na-Ascorbate, 2 Thiourea, 3 Na-Pyruvate, 1 MgSO4, 2 CaCl2; and (3) room temperature (RT) recovery ACSF. Finally, slices were kept at RT in recording ACSF, containing (in mM): 125 NaCl, 26 NaHCO3, 1.25 NaH2PO4, 2.5 KCl, 1 MgCl2, 2 CaCl2, 11 Glucose. During recordings, fresh ACSF was continuously delivered to the slice chamber at ~1.5 ml/min and warmed to 30-32°C with an inline heater (Warner Instruments). Where indicated, the following drugs were added to the recording ACSF: D-AP5 (50 μM, Cayman Chemical), NBQX disodium (5 μM, Tocris Bioscience), TTX (1 μM, Tocris Bioscience), 4-AP (100 μM, Tocris Bioscience), and GBZ (10 μM, Tocris Bioscience). For whole-cell recordings, a high chloride internal solution was used, adjusted to 290±5 mOsm and pH 7.3-7.4, containing (in mM): 130 CsCl, 1 EGTA, 10 HEPES, 5 QX-314-Cl, 10 TEA-Cl, 2 Mg-ATP, 0.3 Na-GTP. For loose seal recordings, a HEPES-buffered synthetic interstitial fluid solution (SIF) was used as internal solution, adjusted to 300±5 mOsm and pH 7.3-7.4, containing (in mM): 140 NaCl, 23 Glucose, 15 HEPES, 3 KCl, 1.5 MgCl2, 1.6 CaCl2. Patch pipettes (3-5 MΩ) were pulled (Narishige, PC-100) from borosilicate glass (Warner Instruments, G150TF-4) and moved with the assistance of a micromanipulator (Sensapex). Cells were visualized with a 40x water-immersion objective (NA 0.8, Olympus, #N2667700) on a microscope (Olympus, BX51WI) equipped with infrared-differential interference imaging (DIC) and a camera (QImaging, Retiga Electro Monochrome). An LED light source (CoolLED, pE-300^white^) was used to illuminate the slice through the objective for targeted patching and for optogenetic stimulation. With the aid of a power meter (Thor Labs, PM130D), the LED power was adjusted to deliver ~20 mW/mm^2^ to the slice during the o-stim. Signals were recorded at 10 kHz using Wavesurfer v0.945 (https://wavesurfer.janelia.org/), a National Instruments digitizer and a Multiclamp 700B amplifier (Molecular Devices). Data analysis was performed offline using custom-written MATLAB scripts.

#### Synaptic connectivity

Bead-labeled cells were held at −70 mV and exposed to the o-stim (5 ms blue light pulse) in 5-10 sweeps, with a 30 s interval between sweeps. Series resistance (Rs) was monitored, but not compensated. Liquid junction potential was not corrected. Cells with Rs > 25 MΩ or with more than 30% change in Rs during the recording were excluded from the dataset. oIPSCs were characterized as fast-onset events (a monotonic decrease in current for 1.5 ms) that happened within 20 ms of the start of the light pulse. In rare sweeps, mIPSCs were mislabeled as oIPSCs. Thus, a *cell* was labeled as “shows an oIPSC” only if oIPSCs were detected in more than 50% of the recorded sweeps. Cells that did not fit this criteria were labeled as “no oIPSC”. For a subset of cells that showed an oIPSC, GBZ was added to the bath for 4 min, and the response to the o-stim was reassessed. Before testing a new cell, GBZ was washed off for at least 20 min. These wash-in and wash-off times were sufficient to block and unblock mIPSCs, respectively (data not shown). The oIPSC amplitude and onset latency reported for each cell were averaged across sweeps. In experiments using the VGAT-IRES-Cre line, a total of 17 cells (4 DLS-projecting and 13 DMS-projecting) were excluded from the dataset due to ChR2 expression, as evidenced by GBZ-insensitive oIPSCs with onset latency < 1 ms. A VGAT+ subgroup of dopaminergic neurons has been previously described (Poulin et al., 2020).

#### Effects on tonic firing

Bead-labeled cells were recorded in voltage clamp (no holding voltage was applied) with a loose seal (20-100 MΩ) and exposed to the o-stim (5 ms pulses delivered at 20 Hz for 3 s) in 5-10 sweeps, with a 30 s interval between sweeps. 10 sweeps were recorded for 88% of the cells (126/142 cells). Cells that did not display tonic firing were excluded from the dataset. The baseline firing rate was calculated during the 3 s prior to the o-stim. Mean±SD were calculated across sweeps. In our experiments using the VGAT-IRES-Cre line, a total of 16 cells (7 DLS-projecting and 9 DMS-projecting) were excluded from the dataset due to ChR2 expression, as evidenced by GBZ-insensitive light-evoked excitation.

#### Approximate cell location

Following each cell recording, a low magnification DIC image was taken with a 5x air-immersion objective (NA 0.15, Olympus, #N2181500) to show the relative position of the cell in the slice. Offline, images from the same slice were stitched in MATLAB for registration purposes. Stitched DIC images were later aligned to an MRI based atlas (Chon et al., 2019) in Adobe Illustrator.

### Histology

#### Slices used exclusively for histology (Figure and Extended Figure 3)

Following Euthasol injection, unresponsive mice were transcardially perfused with ice-cold phosphate-buffered saline (PBS), followed by 4% paraformaldehyde (PFA) diluted in PBS. Brains were immersed in 4% PFA overnight, and then cryoprotected with 30% sucrose (diluted in PBS) at 4°C. Coronal slices (30-50 μm thick) were made using a freezing microtome (Leica, SM2010 R). Staining was performed on free floating slices, with 3×10 min PBS washes in-between incubations. Slices were blocked for 1-2 h at RT with 3% normal goat serum (NGS) diluted in 0.3% PBST (0.3% Triton X-100 in PBS). Then, slices were incubated overnight at 4°C with primary antibodies diluted in blocking solution. Striatum slices were incubated with guinea pig anti-Cre (1:500, Synaptic Systems, #257004) and rabbit anti-GFP (1:1000, Invitrogen, #A11122), while midbrain slices were incubated with chicken anti-TH (1:500, Aves Labs, #TYH) and rabbit anti-GFP (1:1000, Invitrogen, #A11122). Afterwards, slices were incubated for 2-3 h at RT in secondary antibodies diluted in a modified blocking solution (1% NGS in 0.3% PBST). Striatum slices were incubated with goat anti-guinea pig 647 (1:500, Invitrogen, #A21450) and goat anti-rabbit 594 (1:500, Invitrogen, #A11012), while midbrain slices were incubated with goat anti-chicken 647 (1:500, Invitrogen, #A21449) and donkey anti-rabbit 488 (1:500, Jackson Immuno Research, #711-546-152). Striatum slices were further stained for 1-2 h at RT with NeuroTrace 435/455 (1:100 diluted in PBS, Invitrogen, #N21479), a fluorescent Nissl staining. Fluoromount-G (Southern Biotech) was used as mounting media. Slides were imaged with an air-immersion 10x objective (NA 0.45, Nikon, #MRD70105) on an epifluorescence microscope (Keyence, BZ-X800).

#### Slices from electrophysiology experiments (Figures and Extended Figures 1-2; 4-7)

Slices were fixed overnight at 4°C in 4% PFA and stored in PBS at 4°C. Staining was performed on free floating slices as described above, with some modifications – 0.3% PBST was replaced by 0.5% PBST, 10% NGS was used for blocking, and 1% NGS was used to dilute antibodies. Cre staining was performed in striatum slices using guinea pig anti-Cre and goat anti-guinea pig 647. TH staining was performed in midbrain slices using chicken anti-TH and goat anti-chicken 647. EYFP signal was enhanced in all slices with GFP immunolabeling, using rabbit anti-GFP and donkey anti-rabbit 488. Retrobeads did not require enhancement. A custom look-up table was applied in ImageJ to match our colorblind safe color-coding (Wong, 2011). For qualitative visualization of midbrain slices, we adjusted the brightness and contrast of the retrobeads and EYFP channel separately due to the brighter fluorescence of the beads. Analysis of injection spread in DMS/DLS was performed in ImageJ, using the following tools: threshold, median filter, and binary outline. A lower threshold was used for outlining the spread of retrobeads due to their brighter fluorescence in comparison to Cre immunolabeling, but the same analysis parameters were used for all mice. Images were aligned to two striatum sections from the Mouse Brain Atlas (Franklin and Paxinos, 2008), and injection outlines were superimposed in Adobe Illustrator.

### Statistical analyses

Statistical analyses were performed in MATLAB using the Kruskal-Wallis test, a non-parametric version of one-way ANOVA, followed by a Multiple Comparison Test.

## ACKNOWLEDGMENTS

We thank I.M. Raman, M. Bevan, D.J. Surmeier, R. Awatramani, and members of the Lerner laboratory for helpful discussions and critical feedback on the manuscript. We thank G. Palissery and S. Pawelko for assistance with mouse breeding. We thank R. Awatramani for providing the TH-2A-Flpo mouse line. We thank V. Ambrosi for helpful discussions about data analysis in MATLAB. This work was supported by an NIH K99/R00 Award (R00MH109569) to T.N.L. and a NIH-NINDS T32 Award (T32NS041234) to P.A.

## AUTHOR CONTRIBUTIONS

Conceptualization, Methodology, Validation, and Project Administration, P.A. and T.N.L.; Investigation, Formal Analysis, Data Curation and Visualization, and Writing - Original Draft, P.A.; Resources, Writing - Review & Editing, Supervision, and Funding Acquisition, T.N.L.

## EXTENDED FIGURES AND LEGENDS

**Extended Figure 1.**
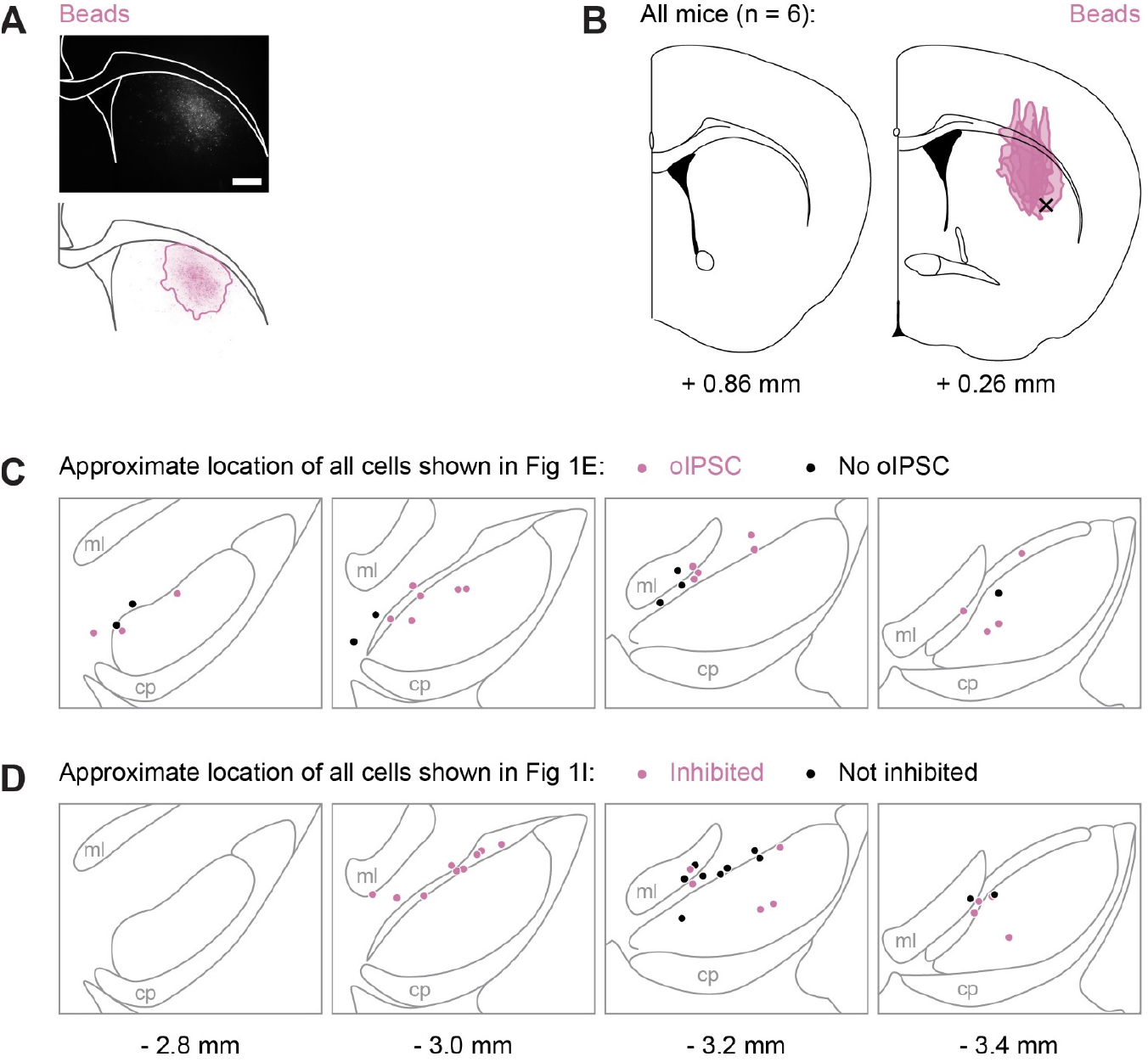
Injection spread in striatum and location of patched cells in midbrain slices. **(A)** Example striatum slice. Scale bar: 0.5 mm. **(B)** Approximate spread of retrobeads in the striatum of all mice used for Figure 1. A black x marks the approximate target location for DLS injections. The numbers below the atlas images indicate their AP position relative to bregma. **(C)** Approximate location of all DLS-projecting cells recorded in whole-cell mode used for Figure 1. Each dot is a cell, color-coded in magenta (oIPSC) or black (no oIPSC). **(D)** Approximate location of all DLS-projecting cells recorded in loose seal mode used for Figure 1. Each dot is a cell, color-coded in magenta (inhibited) or black (not inhibited). The numbers below the atlas images indicate their AP position relative to bregma.

**Extended Figure 2.**
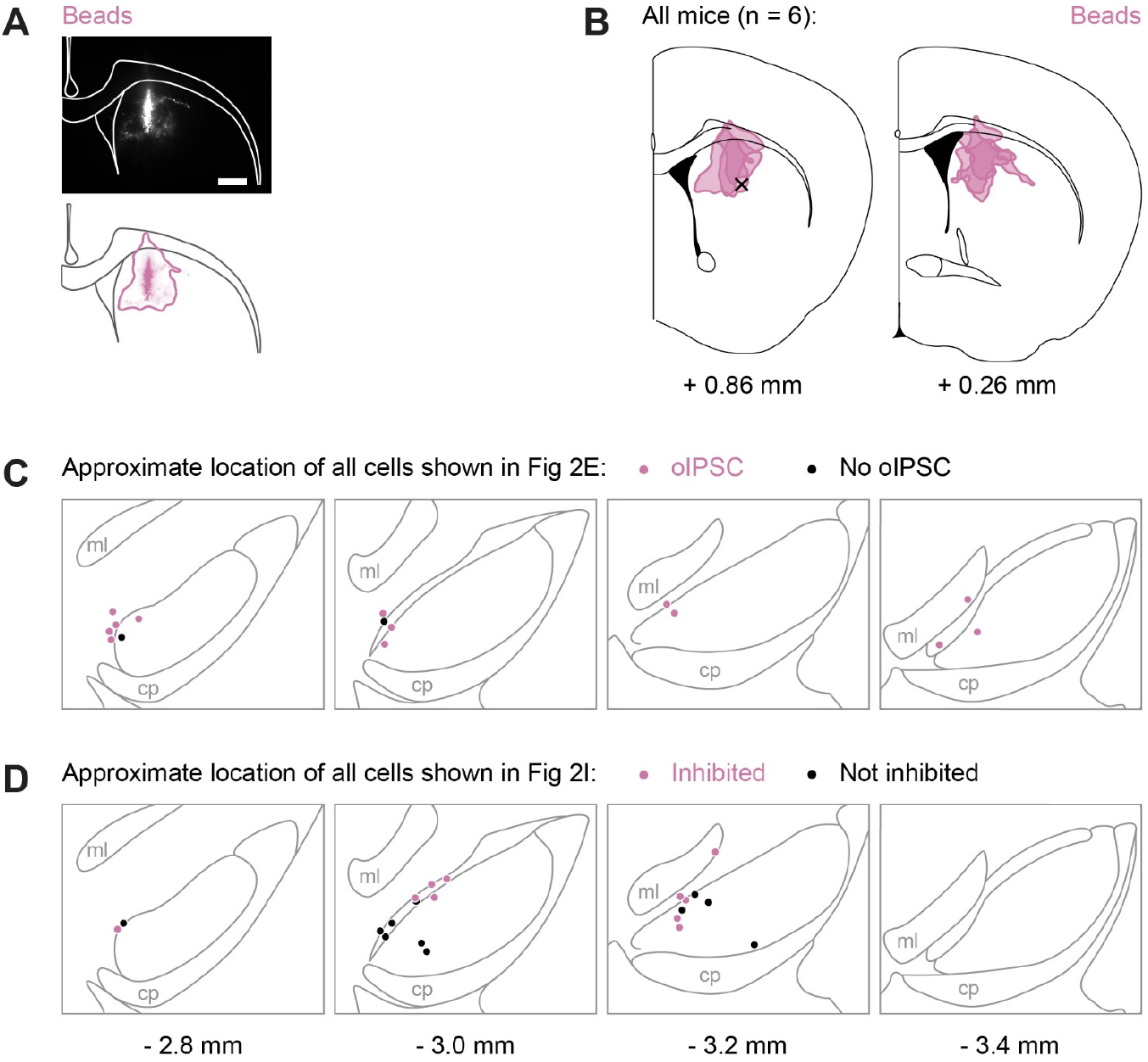
Injection spread in striatum and location of patched cells in midbrain slices. **(A)** Example striatum slice. Scale bar: 0.5 mm. **(B)** Approximate spread of retrobeads in the striatum of all mice used for Figure 2. A black x marks the approximate target location for DMS injections. The numbers below the atlas images indicate their AP position relative to bregma. **(C)** Approximate location of all DMS-projecting cells recorded in whole-cell mode used for Figure 2. Each dot is a cell, color-coded in magenta (oIPSC) or black (no oIPSC). **(D)** Approximate location of all DMS-projecting cells recorded in loose seal mode used for Figure 2. Each dot is a cell, color-coded in magenta (inhibited) or black (not inhibited). The numbers below the atlas images indicate their AP position relative to bregma.

**Extended Figure 3.**
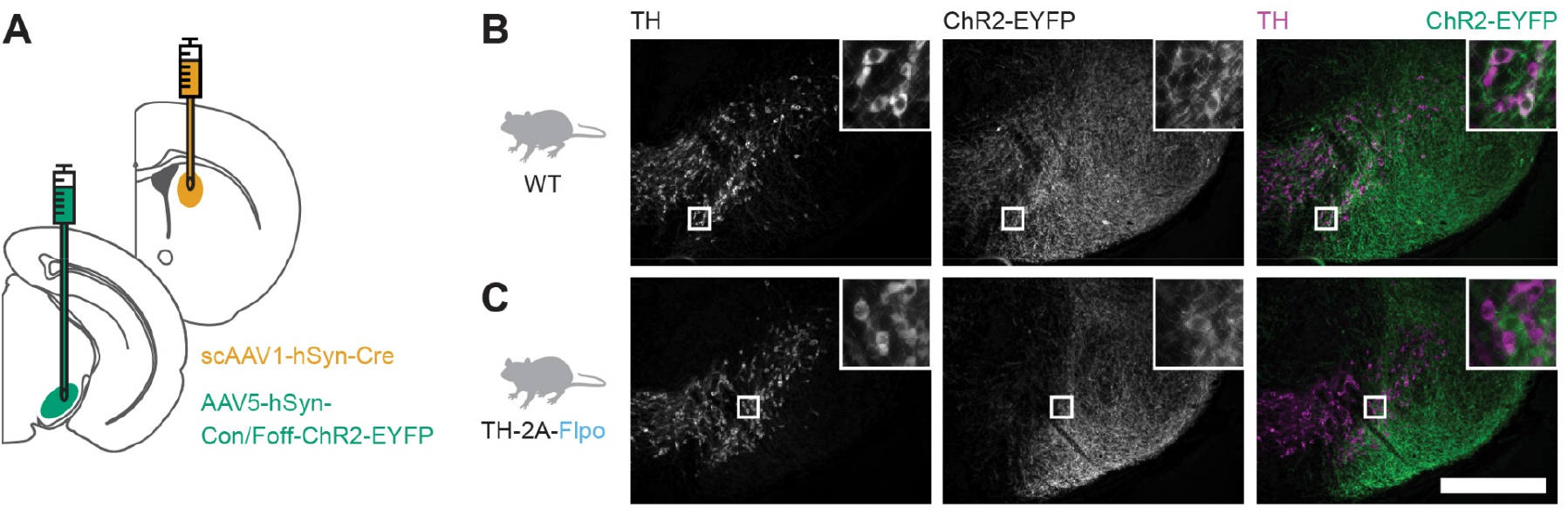
Similar results from Figure 3E are obtained using a Con/Foff-ChR2-EYFP virus. **(A)** Experimental design for labeling DMS-targeted, non-dopaminergic neurons in SNr with ChR2-EYFP. **(B-C)** Example SN histology after injections in WT (B) and TH-2A-Flpo (C) mice. Scale bar: 0.5 mm.

**Extended Figure 4.**
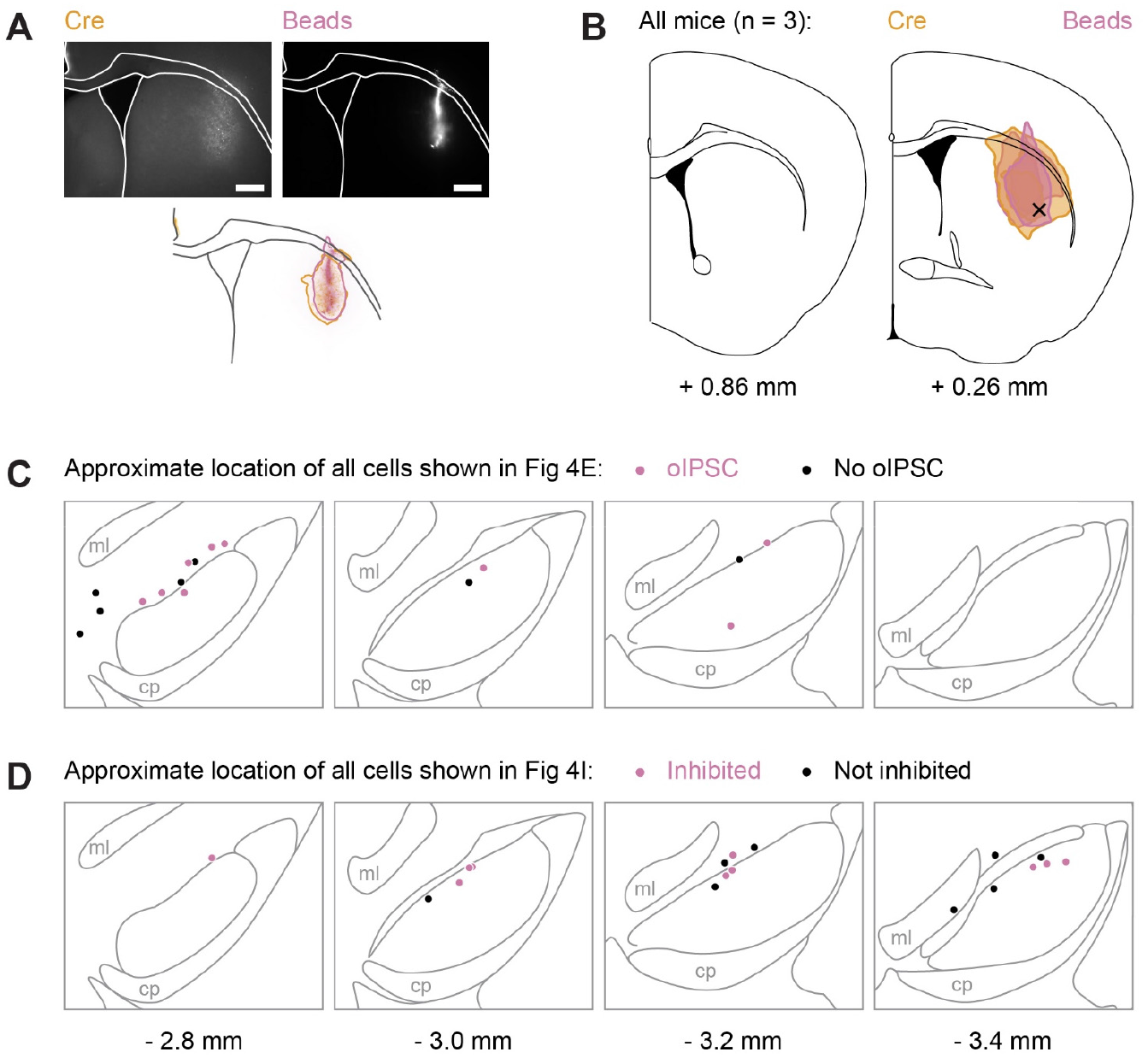
Injection spread in striatum and location of patched cells in midbrain slices. **(A)** Example striatum slice. Scale bar: 0.5 mm. **(B)** Approximate spread of retrobeads in the striatum of all mice used for Figure 4. A black x marks the approximate target location for DLS injections. The numbers below the atlas images indicate their AP position relative to bregma. **(C)** Approximate location of all DLS-projecting cells recorded in whole-cell mode used for Figure 4. Each dot is a cell, color-coded in magenta (oIPSC) or black (no oIPSC). **(D)** Approximate location of all DLS-projecting cells recorded in loose seal mode used for Figure 4. Each dot is a cell, color-coded in magenta (inhibited) or black (not inhibited). The numbers below the atlas images indicate their AP position relative to bregma.

**Extended Figure 5.**
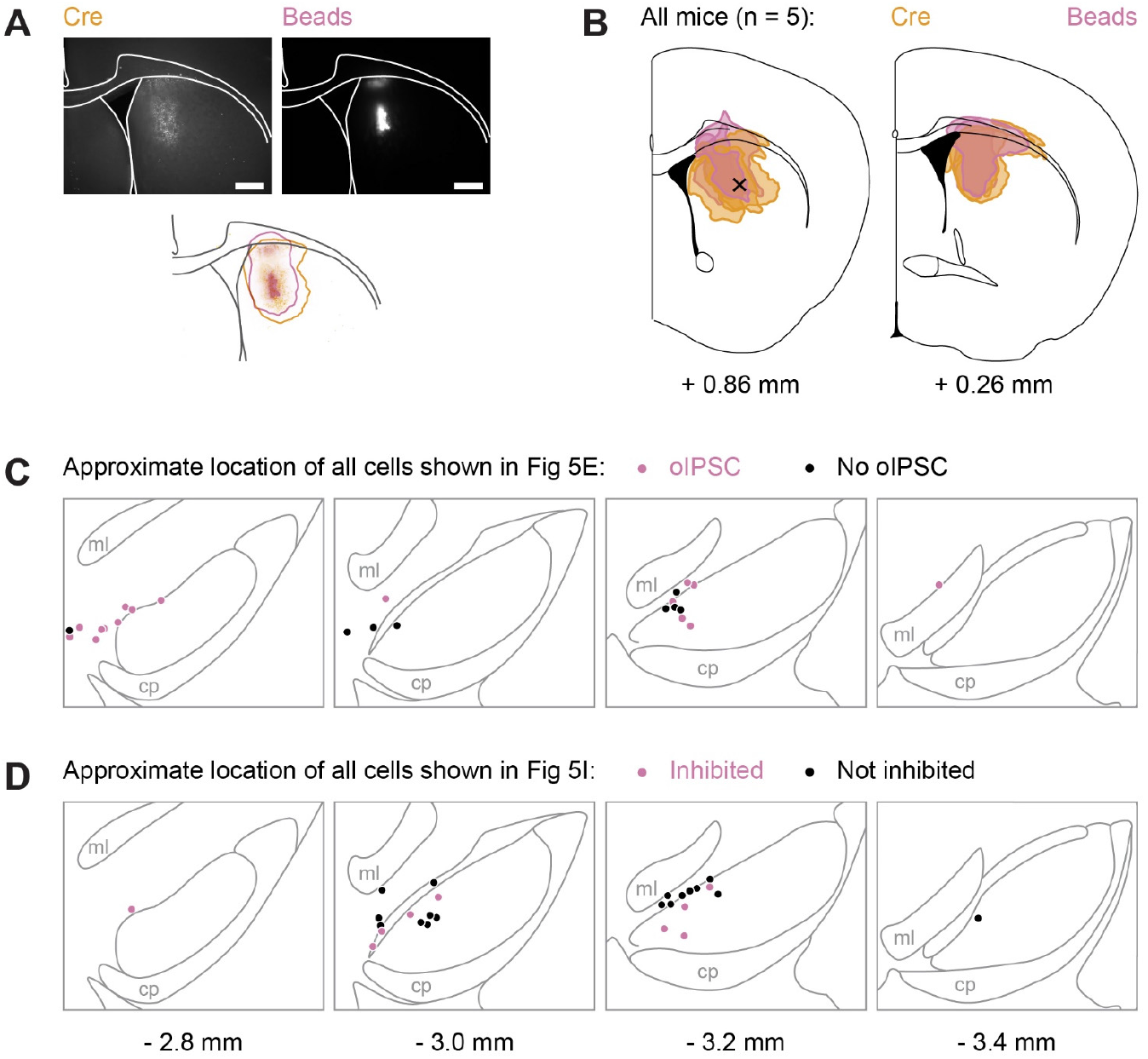
Injection spread in striatum and location of patched cells in midbrain slices. **(A)** Example striatum slice. Scale bar: 0.5 mm. **(B)** Approximate spread of retrobeads in the striatum of all mice used for Figure 5. A black x marks the approximate target location for DMS injections. The numbers below the atlas images indicate their AP position relative to bregma. **(C)** Approximate location of all DMS-projecting cells recorded in whole-cell mode used for Figure 5. Each dot is a cell, color-coded in magenta (oIPSC) or black (no oIPSC). **(D)** Approximate location of all DMS-projecting cells recorded in loose seal mode used for Figure 5. Each dot is a cell, color-coded in magenta (inhibited) or black (not inhibited). The numbers below the atlas images indicate their AP position relative to bregma.

**Extended Figure 6.**
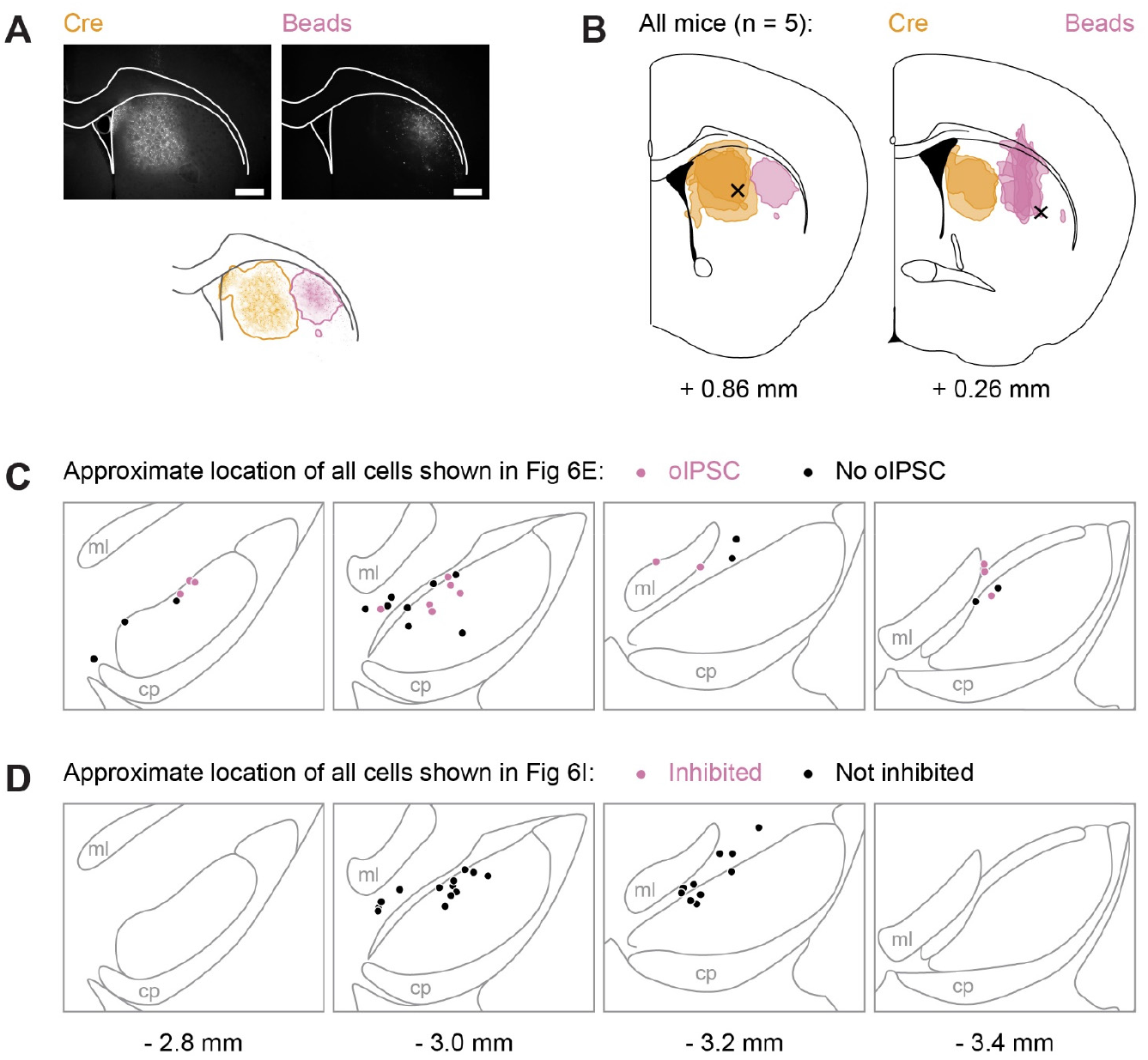
Injection spread in striatum and location of patched cells in midbrain slices. **(A)** Example striatum slice. Scale bar: 0.5 mm. **(B)** Approximate spread of retrobeads in the striatum of all mice used for Figure 6. A black × marks the approximate target location for DMS (left) and DLS (right) injections. The numbers below the atlas images indicate their AP position relative to bregma. **(C)** Approximate location of all DLS-projecting cells recorded in whole-cell mode used for Figure 6. Each dot is a cell, color-coded in magenta (oIPSC) or black (no oIPSC). **(D)** Approximate location of all DLS-projecting cells recorded in loose seal mode used for Figure 6. Each dot is a cell, color-coded in magenta (inhibited) or black (not inhibited). The numbers below the atlas images indicate their AP position relative to bregma.

**Extended Figure 7.**
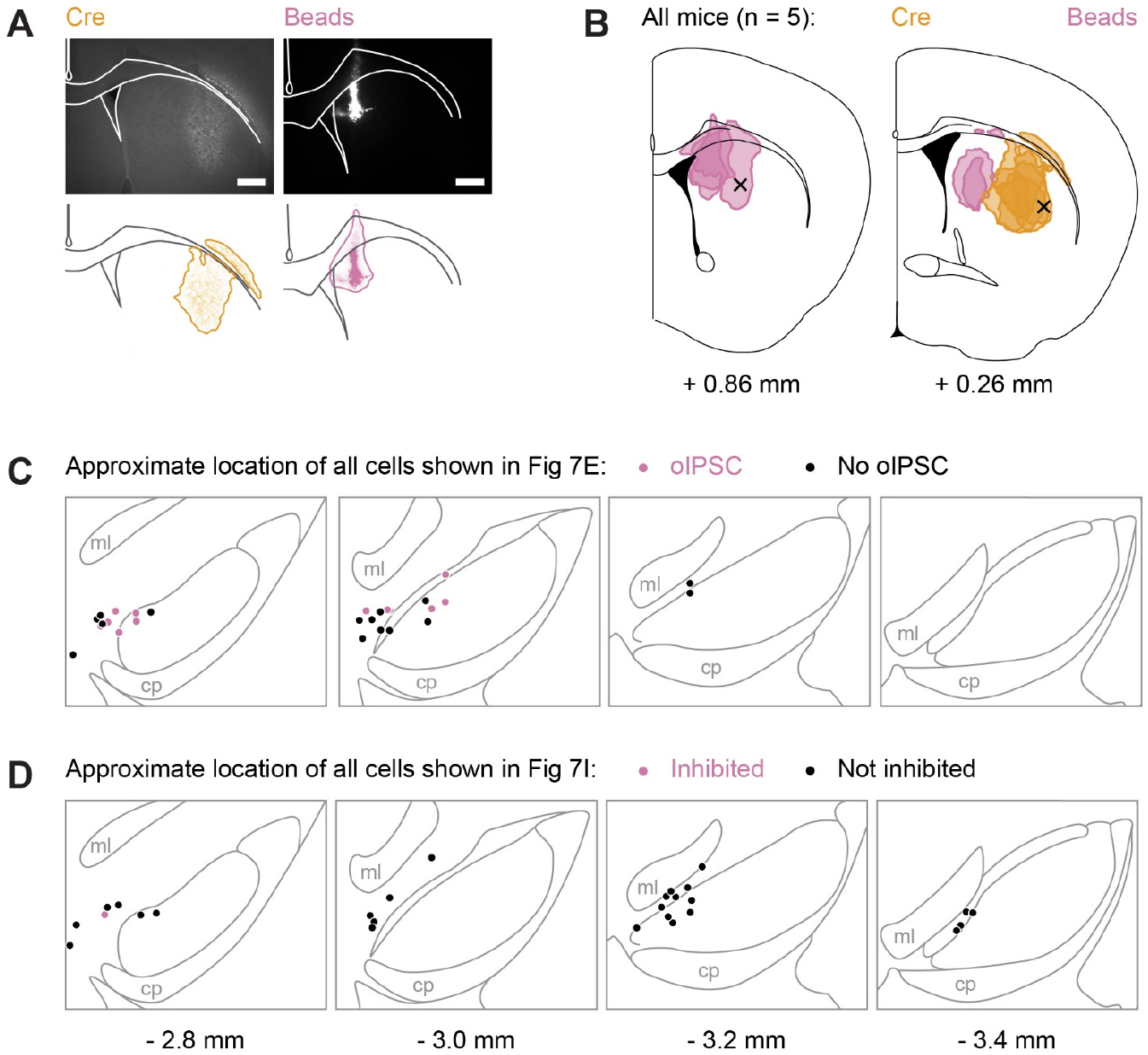
Injection spread in striatum and location of patched cells in midbrain slices. **(A)** Example striatum slice. Scale bar: 0.5 mm. **(B)** Approximate spread of retrobeads in the striatum of all mice used for Figure 7. A black × marks the approximate target location for DMS (left) and DLS (right) injections. The numbers below the atlas images indicate their AP position relative to bregma. **(C)** Approximate location of all DMS-projecting cells recorded in whole-cell mode used for Figure 7. Each dot is a cell, color-coded in magenta (oIPSC) or black (no oIPSC). **(D)** Approximate location of all DMS-projecting cells recorded in loose seal mode used for Figure 7. Each dot is a cell, color-coded in magenta (inhibited) or black (not inhibited). The numbers below the atlas images indicate their AP position relative to bregma.

**Extended Figure 8.**
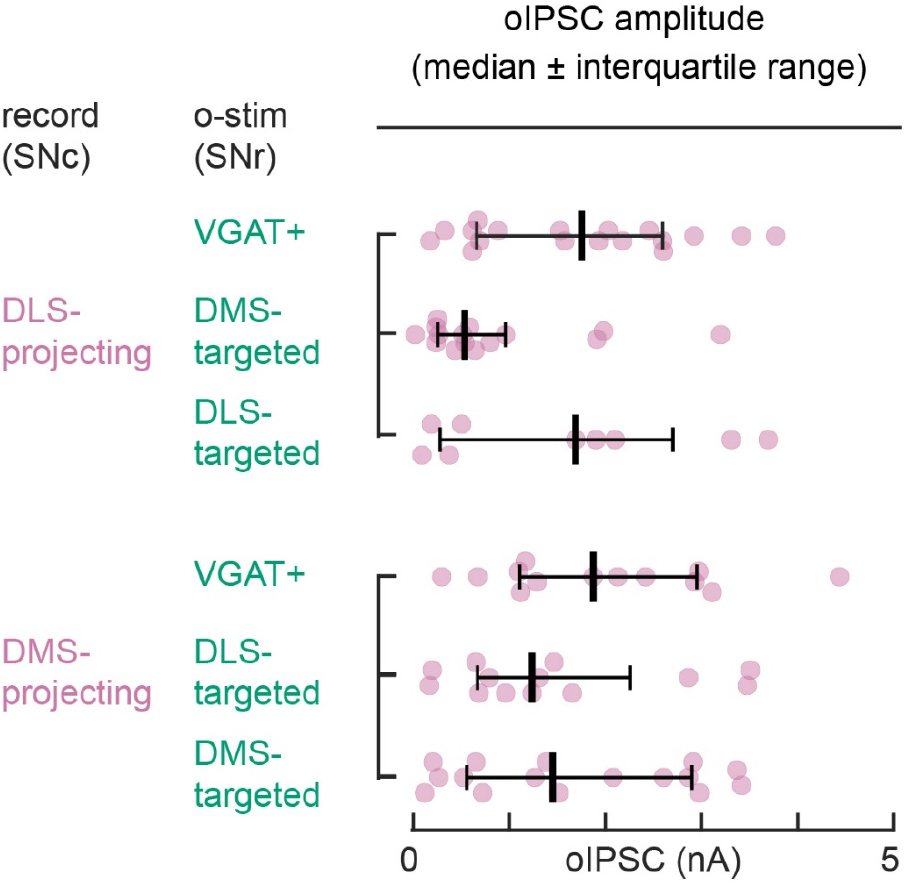
oIPSC amplitude for all tested striato-nigro-striatal circuits. Data from individual cells is shown, along with median and interquartile ranges (adapted from Figures 1-2G and 4-7G). There are no significant differences between groups (Kruskal-Wallis test, p=0.10).

## REFERENCES

Alexander, G.E., DeLong, M.R., and Strick, P.L. (1986). Parallel Organization of Functionally Segregated Circuits Linking Basal Ganglia and Cortex. Annu. Rev. Neurosci. 9, 357–381.

Aoki, S., Smith, J.B., Li, H., Yan, X., Igarashi, M., Coulon, P., Wickens, J.R., Ruigrok, T.J., and Jin, X. (2019). An open cortico-basal ganglia loop allows limbic control over motor output via the nigrothalamic pathway. ELife 8, e49995.

Belin, D., and Everitt, B.J. (2008). Cocaine Seeking Habits Depend upon Dopamine-Dependent Serial Connectivity Linking the Ventral with the Dorsal Striatum. Neuron 57, 432–441.

Brown, H.D., McCutcheon, J.E., Cone, J.J., Ragozzino, M.E., and Roitman, M.F. (2011). Primary food reward and reward-predictive stimuli evoke different patterns of phasic dopamine signaling throughout the striatum. Eur. J. Neurosci. 34, 1997–2006.

Burke, D.A., Rotstein, H.G., and Alvarez, V.A. (2017). Striatal Local Circuitry: A New Framework for Lateral Inhibition. Neuron 96, 267–284.

Cai, Y., and Ford, C.P. (2018). Dopamine Cells Differentially Regulate Striatal Cholinergic Transmission across Regions through Corelease of Dopamine and Glutamate. Cell Rep. 25, 3148–3157.e3.

Chevalier, G., Vacher, S., Deniau, J.M., and Desban, M. (1985). Disinhibition as a basic process in the expression of striatal functions. I. The striato-nigral influence on tecto-spinal/tecto-diencephalic neurons. Brain Res. 334, 215–226.

Corbit, L.H., Nie, H., and Janak, P.H. (2012). Habitual alcohol seeking: time course and the contribution of subregions of the dorsal striatum. Biol. Psychiatry 72, 389–395.

Derusso, A.L., Fan, D., Gupta, J., Shelest, O., Costa, R.M., and Yin, H.H. (2010). Instrumental uncertainty as a determinant of behavior under interval schedules of reinforcement. Front. Integr. Neurosci. 4.

Dorst, M.C., Tokarska, A., Zhou, M., Lee, K., Stagkourakis, S., Broberger, C., Masmanidis, S., and Silberberg, G. (2020). Polysynaptic inhibition between striatal cholinergic interneurons shapes their network activity patterns in a dopamine-dependent manner. Nat. Commun. 11, 5113.

Evans, R.C., Zhu, M., and Khaliq, Z.M. (2017). Dopamine Inhibition Differentially Controls Excitability of Substantia Nigra Dopamine Neuron Subpopulations through T-Type Calcium Channels. J. Neurosci. 37, 3704–3720.

Evans, R.C., Twedell, E.L., Zhu, M., Ascencio, J., Zhang, R., and Khaliq, Z.M. (2020). Functional Dissection of Basal Ganglia Inhibitory Inputs onto Substantia Nigra Dopaminergic Neurons. Cell Rep. 32, 108156.

Farassat, N., Costa, K.M., Stojanovic, S., Albert, S., Kovacheva, L., Shin, J., Egger, R., Somayaji, M., Duvarci, S., Schneider, G., et al. (2019). In vivo functional diversity of midbrain dopamine neurons within identified axonal projections. ELife 8, e48408.

Faure, A., Haberland, U., Condé, F., and Massioui, N.E. (2005). Lesion to the Nigrostriatal Dopamine System Disrupts Stimulus-Response Habit Formation. J. Neurosci. 25, 2771–2780.

Fenno, L.E., Mattis, J., Ramakrishnan, C., Hyun, M., Lee, S.Y., He, M., Tucciarone, J., Selimbeyoglu, A., Berndt, A., Grosenick, L., et al. (2014). Targeting cells with single vectors using multiple-feature Boolean logic. Nat. Methods 11, 763–772.

Fino, E., Vandecasteele, M., Perez, S., Saudou, F., and Venance, L. (2018). Region-specific and state? dependent action of striatal GABAergic interneurons. Nat. Commun. 9, 3339.

Freeze, B.S., Kravitz, A.V., Hammack, N., Berke, J.D., and Kreitzer, A.C. (2013). Control of Basal Ganglia Output by Direct and Indirect Pathway Projection Neurons. J. Neurosci. 33, 18531–18539.

Gerfen, C.R., Herkenham, M., and Thibault, J. (1987). The neostriatal mosaic: II. Patch? and matrix? directed mesostriatal dopaminergic and non-dopaminergic systems. J. Neurosci. 7, 3915–3934.

Grace, A.A., and Bunney, B.S. (1983). Intracellular and extracellular electrophysiology of nigral dopaminergic neurons-1. Identification and characterization. Neuroscience 10, 301–315.

Gremel, C.M., and Costa, R.M. (2013). Orbitofrontal and striatal circuits dynamically encode the shift between goal-directed and habitual actions. Nat. Commun. 4, 2264.

Gulácsi, A., Lee, C.R., Sík, A., Viitanen, T., Kaila, K., Tepper, J.M., and Freund, T.F. (2003). Cell Type? Specific Differences in Chloride-Regulatory Mechanisms and GABAA Receptor-Mediated Inhibition in Rat Substantia Nigra. J. Neurosci. 23, 8237–8246.

Haber, S.N., Fudge, J.L., and McFarland, N.R. (2000). Striatonigrostriatal Pathways in Primates Form an Ascending Spiral from the Shell to the Dorsolateral Striatum. J. Neurosci. 20, 2369–2382.

Hamid, A.A., Frank, M.J., and Moore, C.I. (2021). Wave-like dopamine dynamics as a mechanism for spatiotemporal credit assignment. Cell 184, 2733–2749.e16.

Holly, E.N., Davatolhagh, M.F., Choi, K., Alabi, O.O., Vargas Cifuentes, L., and Fuccillo, M.V. (2019). Striatal Low-Threshold Spiking Interneurons Regulate Goal-Directed Learning. Neuron 103, 92–101.e6.

Ii, E.R.H., Kadoya, K., Hirsch, M., Samulski, R.J., and Tuszynski, M.H. (2008). Efficient Retrograde Neuronal Transduction Utilizing Self-complementary AAV1. Mol. Ther. 16, 296–301.

Ikemoto, S. (2007). Dopamine reward circuitry: Two projection systems from the ventral midbrain to the nucleus accumbens-olfactory tubercle complex. Brain Res. Rev. 56, 27–78.

Joel, D., and Weiner, I. (2000). The connections of the dopaminergic system with the striatum in rats and primates: an analysis with respect to the functional and compartmental organization of the striatum. Neuroscience 96, 451–474.

Khakh, B.S. (2019). Astrocyte-Neuron Interactions in the Striatum: Insights on Identity, Form, and Function. Trends Neurosci. 42, 617–630.

Khirug, S., Yamada, J., Afzalov, R., Voipio, J., Khiroug, L., and Kaila, K. (2008). GABAergic Depolarization of the Axon Initial Segment in Cortical Principal Neurons Is Caused by the Na-K?2Cl Cotransporter NKCC1. J. Neurosci. 28, 4635–4639.

Kramer, P.F., Twedell, E.L., Shin, J.H., Zhang, R., and Khaliq, Z.M. (2020). Axonal mechanisms mediating γ-aminobutyric acid receptor type A (GABA-A) inhibition of striatal dopamine release. ELife 9, e55729.

Lammel, S., Hetzel, A., Häckel, O., Jones, I., Liss, B., and Roeper, J. (2008). Unique Properties of Mesoprefrontal Neurons within a Dual Mesocorticolimbic Dopamine System. Neuron 57, 760–773.

Lammel, S., Ion, D.I., Roeper, J., and Malenka, R.C. (2011). Projection-Specific Modulation of Dopamine Neuron Synapses by Aversive and Rewarding Stimuli. Neuron 70, 855–862.

Lee, J., Wang, W., and Sabatini, B.L. (2020). Anatomically segregated basal ganglia pathways allow parallel behavioral modulation. Nat. Neurosci. 23, 1388–1398.

Lerner, T.N. (2020). Interfacing behavioral and neural circuit models for habit formation. J. Neurosci. Res. 98, 1031–1045.

Lerner, T.N., Shilyansky, C., Davidson, T.J., Evans, K.E., Beier, K.T., Zalocusky, K.A., Crow, A.K., Malenka, R.C., Luo, L., Tomer, R., et al. (2015). Intact-Brain Analyses Reveal Distinct Information Carried by SNc Dopamine Subcircuits. Cell 162, 635–647.

Lipton, D.M., Gonzales, B.J., and Citri, A. (2019). Dorsal Striatal Circuits for Habits, Compulsions and Addictions. Front. Syst. Neurosci. 13.

Liu, C., Goel, P., and Kaeser, P.S. (2021). Spatial and temporal scales of dopamine transmission. Nat. Rev. Neurosci. 22, 345–358.

Lobb, C.J., Troyer, T.W., Wilson, C.J., and Paladini, C.A. (2011). Disinhibition Bursting of Dopaminergic Neurons. Front. Syst. Neurosci. 5.

Lüscher, C., Robbins, T.W., and Everitt, B.J. (2020). The transition to compulsion in addiction. Nat. Rev. Neurosci. 21, 247–263.

Mandelbaum, G., Taranda, J., Haynes, T.M., Hochbaum, D.R., Huang, K.W., Hyun, M., Umadevi Venkataraju, K., Straub, C., Wang, W., Robertson, K., et al. (2019). Distinct Cortical-Thalamic?Striatal Circuits through the Parafascicular Nucleus. Neuron 102, 636–652.e7.

Matsuda, W., Furuta, T., Nakamura, K.C., Hioki, H., Fujiyama, F., Arai, R., and Kaneko, T. (2009). Single Nigrostriatal Dopaminergic Neurons Form Widely Spread and Highly Dense Axonal Arborizations in the Neostriatum. J. Neurosci. 29, 444–453.

Maurin, Y., Banrezes, B., Menetrey, A., Mailly, P., and Deniau, J.M. (1999). Three-dimensional distribution of nigrostriatal neurons in the rat: relation to the topography of striatonigral projections. Neuroscience 91, 891–909.

McElvain, L.E., Chen, Y., Moore, J.D., Brigidi, G.S., Bloodgood, B.L., Lim, B.K., Costa, R.M., and Kleinfeld, D. (2021a). Specific populations of basal ganglia output neurons target distinct brain stem areas while collateralizing throughout the diencephalon. Neuron.

McElvain, L.E., Chen, Y., Moore, J.D., Brigidi, G.S., Bloodgood, B.L., Lim, B.K., Costa, R.M., and Kleinfeld, D. (2021b). Specific populations of basal ganglia output neurons target distinct brain stem areas while collateralizing throughout the diencephalon. Neuron.

Menegas, W., Bergan, J.F., Ogawa, S.K., Isogai, Y., Umadevi Venkataraju, K., Osten, P., Uchida, N., and Watabe-Uchida, M. (2015). Dopamine neurons projecting to the posterior striatum form an anatomically distinct subclass. ELife 4, e10032.

Mohebi, A., Pettibone, J.R., Hamid, A.A., Wong, J.-M.T., Vinson, L.T., Patriarchi, T., Tian, L., Kennedy, R.T., and Berke, J.D. (2019). Dissociable dopamine dynamics for learning and motivation. Nature 570, 65–70.

Petreanu, L., Mao, T., Sternson, S.M., and Svoboda, K. (2009). The subcellular organization of neocortical excitatory connections. Nature 457, 1142–1145.

Poulin, J.-F., Caronia, G., Hofer, C., Cui, Q., Helm, B., Ramakrishnan, C., Chan, C.S., Dombeck, D.A., Deisseroth, K., and Awatramani, R. (2018). Mapping projections of molecularly defined dopamine neuron subtypes using intersectional genetic approaches. Nat. Neurosci. 21, 1260–1271.

Poulin, J.-F., Gaertner, Z., Moreno-Ramos, O.A., and Awatramani, R. (2020). Classification of Midbrain Dopamine Neurons Using Single-Cell Gene Expression Profiling Approaches. Trends Neurosci. 43, 155–169.

Rahmati, N., Normoyle, K.P., Glykys, J., Dzhala, V.I., Lillis, K.P., Kahle, K.T., Raiyyani, R., Jacob, T., and Staley, K.J. (2021). Unique actions of GABA arising from cytoplasmic chloride microdomains. J. Neurosci.

Seiler, J.L., Cosme, C.V., Sherathiya, V.N., Bianco, J.M., and Lerner, T.N. (2020). Dopamine Signaling in the Dorsomedial Striatum Promotes Compulsive Behavior. BioRxiv 2020.03.30.016238x.

Sommer, W.H., Costa, R.M., and Hansson, A.C. (2014). Dopamine systems adaptation during acquisition and consolidation of a skill. Front. Integr. Neurosci. 8, 87.

Tarfa, R.A., Evans, R.C., and Khaliq, Z.M. (2017). Enhanced Sensitivity to Hyperpolarizing Inhibition in Mesoaccumbal Relative to Nigrostriatal Dopamine Neuron Subpopulations. J. Neurosci. 37, 3311–3330.

Tepper, J.M., and Lee, C.R. (2007). GABAergic control of substantia nigra dopaminergic neurons. In Progress in Brain Research, J.M. Tepper, E.D. Abercrombie, and J.P. Bolam, eds. (Elsevier), pp. 189–208.

Tepper, J.M., Martin, L.P., and Anderson, D.R. (1995). GABAA receptor-mediated inhibition of rat substantia nigras dopaminergic neurons by pars reticulata projection neurons. J. Neurosci. 15, 3092–3103.

Thorn, C.A., Atallah, H., Howe, M., and Graybiel, A.M. (2010). Differential Dynamics of Activity Changes in Dorsolateral and Dorsomedial Striatal Loops during Learning. Neuron 66, 781–795.

Tsutsui-Kimura, I., Matsumoto, H., Akiti, K., Yamada, M.M., Uchida, N., and Watabe-Uchida, M. (2020). Distinct temporal difference error signals in dopamine axons in three regions of the striatum in a decision-making task. ELife 9, e62390.

Watabe-Uchida, M., Zhu, L., Ogawa, S.K., Vamanrao, A., and Uchida, N. (2012). Whole-Brain Mapping of Direct Inputs to Midbrain Dopamine Neurons. Neuron 74, 858–873.

Willuhn, I., Burgeno, L.M., Everitt, B.J., and Phillips, P.E.M. (2012). Hierarchical recruitment of phasic dopamine signaling in the striatum during the progression of cocaine use. Proc. Natl. Acad. Sci. 109, 20703–20708.

Xu, M., Kobets, A., Du, J.-C., Lennington, J., Li, L., Banasr, M., Duman, R.S., Vaccarino, F.M., DiLeone, R.J., and Pittenger, C. (2015). Targeted ablation of cholinergic interneurons in the dorsolateral striatum produces behavioral manifestations of Tourette syndrome. Proc. Natl. Acad. Sci. 112, 893–898.

Yang, H., de Jong, J.W., Tak, Y., Peck, J., Bateup, H.S., and Lammel, S. (2018). Nucleus Accumbens Subnuclei Regulate Motivated Behavior via Direct Inhibition and Disinhibition of VTA Dopamine Subpopulations. Neuron 97, 434–449.e4.

Yin, H.H., and Knowlton, B.J. (2006). The role of the basal ganglia in habit formation. Nat. Rev. Neurosci. 7, 464–476.

Yin, H.H., Knowlton, B.J., and Balleine, B.W. (2004). Lesions of dorsolateral striatum preserve outcome expectancy but disrupt habit formation in instrumental learning. Eur. J. Neurosci. 19, 181–189.

Yin, H.H., Knowlton, B.J., and Balleine, B.W. (2005a). Blockade of NMDA receptors in the dorsomedial striatum prevents action-outcome learning in instrumental conditioning. Eur. J. Neurosci. 22, 505–512.

Yin, H.H., Ostlund, S.B., Knowlton, B.J., and Balleine, B.W. (2005b). The role of the dorsomedial striatum in instrumental conditioning. Eur. J. Neurosci. 22, 513–523.

Yin, H.H., Knowlton, B.J., and Balleine, B.W. (2006). Inactivation of dorsolateral striatum enhances sensitivity to changes in the action-outcome contingency in instrumental conditioning. Behav. Brain Res. 166, 189–196.

Yin, H.H., Mulcare, S.P., Hilário, M.R.F., Clouse, E., Holloway, T., Davis, M.I., Hansson, A.C., Lovinger, D.M., and Costa, R.M. (2009). Dynamic reorganization of striatal circuits during the acquisition and consolidation of a skill. Nat. Neurosci. 12, 333–341.

Zingg, B., Chou, X., Zhang, Z., Mesik, L., Liang, F., Tao, H.W., and Zhang, L.I. (2017). AAV-Mediated Anterograde Transsynaptic Tagging: Mapping Corticocollicular Input-Defined Neural Pathways for Defense Behaviors. Neuron 93, 33–47.

Zingg, B., Peng, B., Huang, J., Tao, H.W., and Zhang, L.I. (2020). Synaptic Specificity and Application of Anterograde Transsynaptic AAV for Probing Neural Circuitry. J. Neurosci. 40, 3250–3267.

